# *Wolbachia* infection induces host cell state changes *in vitro* and determines symbiotic fate in *Drosophila*

**DOI:** 10.1101/2025.06.05.658111

**Authors:** Jodie Jacobs, Jonas Nykamp, Alexandra Lum, Cade Mirchandani, William E Seligmann, Samuel Sacco, Merly Escalona, Richard E Green, Shelbi L Russell

## Abstract

Host-associated bacteria influence eukaryotic cellular differentiation, but the underlying mechanisms are difficult to disentangle because of cell-type-specific tropisms and phenotypes. The association between *Wolbachia pipientis* and *Drosophila melanogaster* offers a tractable system for gaining high-resolution insight into how host and microbe gene expression interact. Using *Wolbachia*-infected *D. melanogaster* cell lines, we show that the native *w*Mel strain alters host phenotypic, transcriptomic, and chromatin conformation states. High-throughput imaging reveals that *w*Mel infection differentially alters host cellular phenotypes in a titer-independent manner in some cell lines. These phenotypically altered host cells exhibit differential gene expression complementary to the expression patterns observed in their *w*Mel symbionts. Infection was also negatively associated with chromatin organization and histone modification pathways, prompting us to investigate differences in chromatin conformation between infection states. Micro-C chromatin contact data reveal that *w*Mel infection conferred increased *D. melanogaster* chromatin contacts coinciding with both increased compartmentalization and derepression. The X chromosome exhibited evidence consistent with altered dosage compensation, with no downstream alteration of the X:autosome transcript ratio or X-linked histone acetylation, consistent with a dosage compensation-mediated male-killing mechanism encoded by *w*Mel that has been suppressed in *D. melanogaster*. Overall, these findings suggest that the character of *Wolbachia* symbiosis, cooperative or antagonistic, emerges in response to the host cellular environment. Furthermore, these results demonstrate that *in vitro* cell lines offer promising avenues for uncovering the functional genetic basis of microbe-induced host phenotypes.

## Introduction

The intracellular bacterium *Wolbachia pipientis* is leveraged as a biological control agent to inhibit the reproduction of problematic hosts and suppress the transmission of human diseases (Medina et al. 2019) through the unique host phenotypes it induces (Werren et al. 2008), but little is known about the genetic or mechanistic basis for many of these phenotypes. *Wolbachia* are maternally transmitted through the female germline of their hosts and many strains are capable of manipulating host reproduction to increase the proportion of infected females in the population (Iturbe-Ormaetxe et al. 2011). These bacteria are leveraged as non-native infections in problematic host species, such as *Aedes aegypti* and *Anopheles sp.* (Iturbe-Ormaetxe et al. 2011; Zhang and Dimopoulos 2026), to decrease host population size (Beebe et al. 2021), and to suppress the transmission of arboviral diseases to humans (Utarini et al. 2021). Two host-derived phenotypes underlie these biological control measures: Cytoplasmic incompatibility leverages a germline-based toxin-antidote system to create incompatible mating outcomes between infected males and uninfected females, or females infected with an incompatible strain (Turelli et al. 2022). The ability for some *Wolbachia* strains to block viral transmission is a multifaceted somatic process that revolves around core incompatibilities between the growth requirements of the two intracellular microbes (Mushtaq et al. 2024; Lindsey et al. 2018).

Other host-derived phenotypes exhibited by *Wolbachia* are as diverse as the cell types (Pietri et al. 2016; Frydman et al. 2006; Toomey and Frydman 2014; Toomey et al. 2013) and host species they infect. Other reproductive manipulations have been observed across Arthropoda, from male-killing in Lepidoptera, to feminization in Isopoda, and parthenogenesis in haplodiploid taxa, such as Hymenoptera (Fricke and Lindsey 2024; Werren et al. 2008; Kaur et al. 2021; Iturbe-Ormaetxe and O’Neill 2007; Landmann 2020). Beneficial reproductive phenotypes evolve once infections are fixed in host populations (Turelli 1994). In fruit flies and plant hoppers, *Wolbachia* has been shown to reinforce host fertility through interacting with developmental pathways (Weeks et al. 2007; Russell et al. 2023; Starr and Cline 2002; Flores et al. 2015; Ote et al. 2016) and through nutritional mutualism (Lindsey et al. 2025; Ju et al. 2020). B-vitamin limitation underlies the obligate association between the *w*Cle strain and bed bugs, which rely on *Wolbachia* for this nutrient that is lacking in the host’s diet (Hosokawa et al. 2010; Wiles et al. 2025). In filarial nematodes (Foray et al. 2018; Landmann et al. 2011) and *Asobara tabida* wasps (Dedeine et al. 2001), *Wolbachia* is required for oogenesis. The *w*Bm strain that obligately infects *Brugia malayi* filarial worms and controls germline stem cell quiescence, as well as the mitotic proliferation of differentiating germ cells (Foray et al. 2018), and their loss causes cell death in late oogenesis and early embryogenesis (Landmann et al. 2014). Similarly, the *w*Atab3 strain is essential for *A. tabida* to proceed through the mid-oogenesis checkpoint, and their loss induces cell death (Pannebakker et al. 2007).

Despite the strength of *Wolbachia-*induced host phenotypes, tracking down their genetic underpinnings has been remarkably challenging. Little is known about how *Wolbachia* and host gene expression interact because many studies report minimal or conflicting differential expression patterns due to infection (Mateos et al. 2019; Bennuru et al. 2016; Gutzwiller et al. 2015; Kremer et al. 2012; Darby et al. 2012; Chung et al. 2020). Many strains are distributed broadly across host tissues, but induce tissue-specific phenotypes. This suggests that *Wolbachia’s* host-derived phenotypes may be induced by specific host cell states. In this case, *in vivo* bulk transcriptomic approaches have limited power to identify the underlying differentially expressed genes. Indeed, this may explain why bulk transcriptomic data were insufficient to resolve the genes *Wolbachia* use to rescue loss of fertility in *Drosophila melanogaster* ovaries (Russell et al. 2023). Thus, *Wolbachia* transcriptomics faces challenges with detecting rare and ephemeral cell transcriptomes that recapitulate those in developmental (Tang et al. 2009), neuro– (Lovatt et al. 2012), and cancer biology (Conde-Lopez et al. 2024).

Homogeneous *Wolbachia*-infected *Drosophila* cell cultures enable the interrogation of cell-type-specific microbe-host interactions through isolating and concentrating the effects of infection. We leverage the *w*Mel strain that naturally occurs in *D. melanogaster* and reaches high titers *in vitro*, and has been applied to control mosquito populations in the field. Through high-throughput imaging, dual transcriptome sequencing, and chromatin contact sequencing, we reveal that *w*Mel induces cell type-specific changes in host phenotype and gene expression that are hard-coded into the epigenetic landscape. We find that *Wolbachia* gene expression is differentially impacted by host cell type in a way that complements the effects of infection on host gene expression. In these immortal somatic cell lines, we find preliminary evidence for mutualistic and pathogenic interactions between wMel and *D. melanogaster* that depend on host cell state.

## Results

Here, we show that *Wolbachia* and *Drosophila* gene expression interact to alter bacterial and eukaryotic cell states, and potentially alter the cost-benefit dynamics of infection. To generate strain and passage-matched infected and uninfected cell lines, we infected S2 cells with the *w*Mel variant naturally present in the JW18 cell line, and cured versions of both lines with doxycycline, validating infection states with FISH (described in Methods). We imaged cellular morphology in high-throughput to quantitatively assess the phenotypic impact of infection on two different host cell types. Cell pellets were processed in bulk for dual host and microbe transcriptome sequencing and chromatin contact sequencing (resource outlined in Fig. 1A and Methods outlined in Supplemental Fig. S1, Tables S1-S3). The data presented below suggest that *Wolbachia* responds to the gene expression profile of the host cell, inducing complementary changes in host gene expression. Overall, these results support the use of *in vitro* systems to empower identifying the genetic basis for microbe-derived host phenotypes.

**Figure 1.**
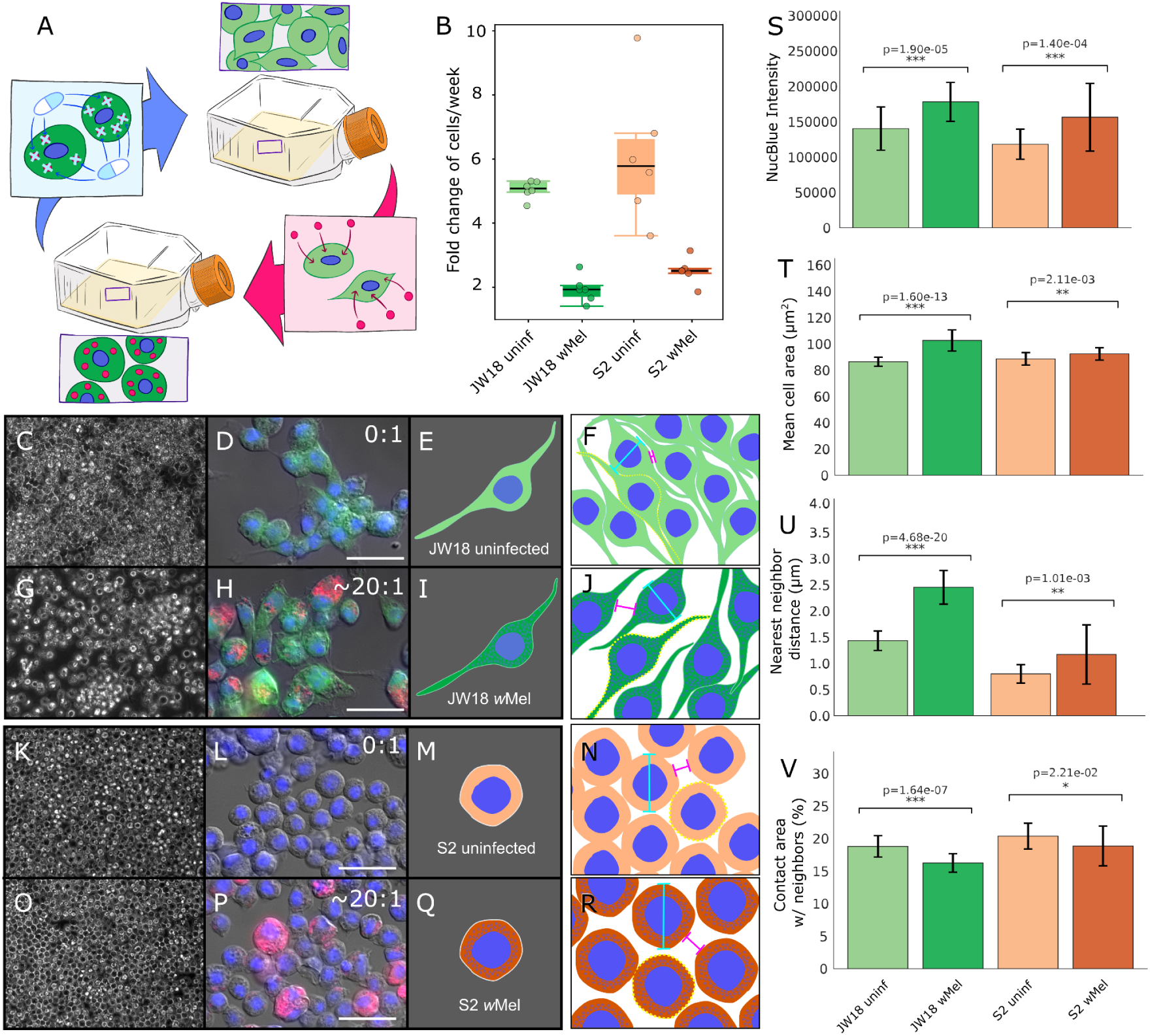
*Wolbachia* infection alters *Drosophila* cell line morphologies. A) Schematic of the shell-vial infection and DOX-curing protocols to make strain and cell line-matched experimental infection systems. B) Cell growth rates quantified by hemocytometer. C-R) Micrographs and illustrated depictions of the immortalized *D. melanogaster* C-J) JW18 and K-R) S2 cell lines. C-F,K-N) uninfected and G-J,O-R) infected with *w*Mel *Wolbachia*. C,G,K,O) Live cultures imaged at 40x on a tissue culture microscope. D,H,L,P) Fixed and stained cells imaged on an epifluorescent compound microscope. Green=Jupiter-GFP, Blue=DAPI, Red=16S rRNA. Scale bars=25 µm. Overlaid ratios indicate the average *w*Mel:D.mel genomic titer measured by Illumina sequencing. C,F,I,L) E,I,M,Q) Schematic representations of the cell lines and their morphologies, colored by cell type and infection state. Green represents JW18 cells and orange represents S2 cells. Lighter shading reflects the uninfected states. Darker shading reflects the *w*Mel-infected cell states. F,J,N,R) Illustrations of the parameters measured on the Phenix imager. S-V) Bar plots of average measurements made with the Phenix imager and analyzed with Harmony. Standard deviations are shown with error bars. Two tailed Welch’s t-test p-values.

### Intracellular *Wolbachia* infection alters host cell morphology and growth

We quantified the phenotypic impacts of infection in high-throughput with a Phenix imager, finding that infection of *D. melanogaster* cells with the *w*Mel strain significantly alters host cell growth rate and morphology (Fig. 1 and Supplemental Fig. S2, Table S1). Quantifying the impact of infection with *w*Mel on host cell growth confirmed that infection slows the growth of both S2 and JW18 cell lines by ⅔-½x compared to the uninfected state (Fig. 1B, as in (Mirchandani et al. 2024)). Visually, the JW18 and S2 cells lines appear distinct with *w*Mel infection altering their morphology (Fig. 1C-R). Next, we used a Phenix imager to quantify *w*Mel infection titer with NucBlue staining, revealing a similar infection intensity between *w*Mel-infected S2 and JW18 cells (Fig. 1S), consistent with genomic titer estimates for these cell lines (Mirchandani et al. 2024). Despite similar titers, JW18 cells increase in mean area more upon infection than S2 cells do (Fig 1T), although both significantly increase in size (103+/-55 µm infected versus 86+/-33 µm uninfected, p=1.60e-13 and 93+/-42 µm infected versus 89+/-41 µm uninfected, p=2.11e-3, respectively, Welch’s t-test). The JW18 cell line displayed increased nearest neighbor distance (2.5+/-4.8 µm infected versus 1.4+/-3.2 µm uninfected, p=4.68e-20 Welch’s t-test) and reduced contact area with neighbors (16.2+/-17.5 µm infected versus 18.8+/− 18.3 µm uninfected, p=1.64e-7 Welch’s t-test) at confluency in the infected compared to uninfected state (Fig. 1U-V). The S2 cell line exhibited a similar, but less pronounced increase in nearest neighbor distance (1.3+/-3.0 µm infected versus 0.80+/-2.10 µm uninfected, p=1.01e-3 Welch’s t-test) and reduced contact area with neighbors (17.5+/-17.3 µm infected versus 20.4+/-18.1 µm uninfected, p=2.21e-2 Welch’s t-test) at confluence in the infected compared to uninfected state (Fig. 1U-V).

### Bulk RNAseq supports and describes the phenotypic distinction between JW18 and S2 cell states

The transcriptomic states of *Drosophila* S2 and JW18 cells with and without *w*Mel infection support and explain the distinction in their uninfected and infected morphologies (Fig. 2 and Supplemental Figs. S3-S4). After extracting and normalizing read pseudoalignments to the host transcriptome, we performed principal component analysis (PCA) to identify the main axes of variation in the data. Tight clustering of samples by cell line and infection state supports our morphological observations that each *D. melanogaster* cell type is highly consistent within a cell line and distinct between cell lines (Fig. 2A and Supplemental Fig. S3E). The top four principal components (PCs) explained more than 98% of transcriptomic count variance (Fig. 2A and Supplemental Fig S3A-D), with PC1 separating cells by type and accounting for 74.73% of variance. Infection with *Wolbachia* explained another 12% of eigenvalue variance in PC2. Many of the top marker genes driving PCA clustering were uncharacterized (Fig. 2B).

**Figure 2.**
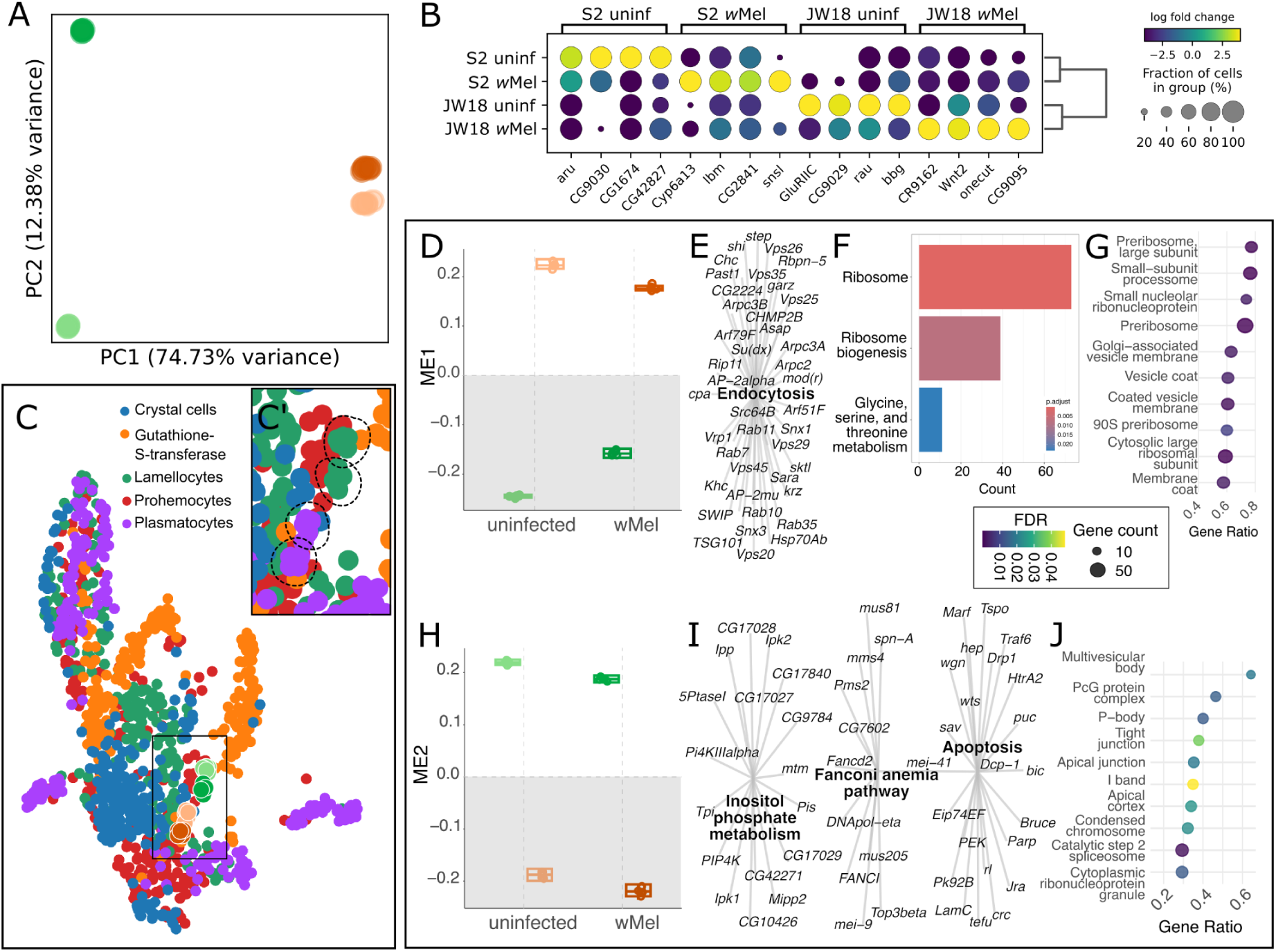
*Wolbachia* infection alters *Drosophila* cell line transcriptomic state, although both retain a hemocyte-like identity due to culture conditions. A) Principal Component Analysis (PCA) revealing distinct transcriptomic profiles by cell line and infection status (orange=JW18, green=S2). B) Heatmap showing the top four differentially expressed marker genes contributing to the PCA by Wilcoxon Rank Sum p-value, with circle size indicating the fraction of cells expressing each gene. C) UMAP projections of bulk RNA-seq data onto the *Drosophila* myeloid-like single-cell reference atlas. C’) Highlights where our bulk cell transcriptomes mapped to the embryo atlas. Dotted outlines of where the bulk samples mapped, with the points removed, so that the alignment of JW18 cells with lamellocyte clusters and S2 with plasmatocyte clusters can be seen. D-J) WGCNA module eigengene evidence associated with the D-G) S2 cell type (module 1) and H-J) JW18 cell type (module 2). D,H) Eigengene value plots for each module, separated by sample type. KEGG enrichment network plots for each module (FDR-adjusted p-value<0.05) from E, I) ShinyGO and F) clusterProfiler. G,J) GO component enrichment plots for each module, plotting the top 5-10 terms by gene ratio (FDR-adjusted p-value<0.05).

We characterized the distinction among cell lines and confirmed their similarity to hemocytes by projecting normalized bulk RNAseq transcript counts onto three Fly Cell Atlases (Fig. 2C and Supplemental Fig. S4). Both cell lines in both infection states mapped to the adult cell atlases (Fig. 2C and Supplemental Fig. S4A), but they failed to map to cell clusters in the embryonic and larval atlas (Supplemental Fig. S4B). This may reflect terminal differentiation of the *in vitro* lines, which have been growing continuously in culture for decades (Serbus et al. 2012a; Schneider 1972). While the cell lines mapped to the adult Fly Cell Atlas, which was produced from “bloodless” tissues, they did not map to well defined clusters or known cell types other than the “body” (Supplemental Fig. S4A). Instead, they mapped closer to known clusters in the blood cell atlas (Fig. 2C). The uninfected and infected versions of the JW18 cell line clustered with lamellocytes, and the two versions of the S2 cell line clustered with plasmatocytes. The hemocyte-like identity of these cell lines is not unexpected because the cell culture media types they were immortalized with were designed to recapitulate the chemical environment of hemolymph (Luhur et al. 2020). While they are clustered closer to blood cell types than any other *in vivo* cell type, they are on the outskirts of those clusters, which may reflect novel gene expression programs induced due to immortalization. Infection state may also impact clustering, given the uninfected statuses of the *Drosophila* cell atlases (Supplemental Fig. S5).

### Transcriptomic analyses reveal host genes correlated with cell type or infection state

We identified thousands of host genes that change in expression due to cell type, infection state, or both through differential expression analysis of the host transcriptome (Supplemental Fig. S6A-C and Tables S4-6). After quality control and processing, we detected 10,839 expressed *D. melanogaster* genes among samples. Cell type corresponded to 5,946 genes with greater than 0.5x log2-fold change across replicates, with 1,948 genes having greater than 2x log2-fold change (p ≤ 0.01 FDR-corrected Wald Test, Table S4). Infection with *w*Mel corresponded to the differential expression of 3,276 genes at 0.5x or greater log2-fold change, and 691 genes at greater than 2x log2-fold change (p ≤ 0.01 FDR-corrected Wald Test), with more genes upregulated in the infected than uninfected state (562 upregulated vs 129 down regulated at 2x log2-fold; Supplemental Fig. S6B and Table S5). The interaction between cell type and infection was highly significant: 4,475 genes were differentially expressed due to the interaction term at 0.5x log2-fold or greater change, and 1,190 genes at 2x log2-fold change (p ≤ 0.01 FDR-corrected Wald Test, Table S6). The high consistency among replicates (Fig. 2A and Supplemental Fig. S6E-G) suggests that these data are sufficiently powered to detect changes in gene expression down to a 1.4x difference (0.5x log2-fold change) in transcript counts between conditions (Schurch et al. 2016).

Gene set enrichment analysis (GSEA) of the differentially expressed *D. melanogaster* genes revealed dozens of functional pathways associated with cell type, infection state, and the interaction between cell type and infection (p ≤ 0.05 permutation test; Supplemental Fig. S6H-J, Tables S7-12). The JW18 cell type was associated with downregulated ribosome expression and Wnt signalling (Supplemental Fig. S6H and Tables S7-8). Compared to the S2 cell type, JW18 cells upregulated their sugar metabolism, cell surface mucopolysaccharide (glycosaminoglycan) biosynthesis, DNA replication and repair, and the polycomb repressive complex chromatin modifiers. Infection was associated with the downregulation of bacterial invasion response and fatty acid degradation genes, and the upregulation of purine metabolism (Supplemental Fig. S6I and Tables S9-10). Several of these functions overlapped the gene set enriched for interactions between cell type and infection (Supplemental Fig. S6J and Tables S11-12), suggesting that some pathways jointly interacted with infection and cell type.

Using weighted gene co-expression network analysis (WCGNA), we inferred 13 clusters of *D. melanogaster* genes with correlated expression patterns, termed modules, 11 of which were significantly associated with cell type, infection, or both (p ≤ 0.05, Benjamini–Hochberg FDR-corrected moderated t-test, Supplemental Fig. S7-12, Fig. 2D-J, Fig. 3A, B-E, I-J, O-Q, T-X, and Tables S13-15). An eigengene value, or module eigengene (ME), is the first component from the PCA of the gene expression matrix for the module that explains the largest variance in the module’s overall expression and acts as a weighted summary of the expression profile (Langfelder and Horvath 2007). Nearly all of the genes in the significant modules were also differentially expressed (89.0%-99.5%, Supplemental Fig. S9A). The distinction between S2 and JW18 cell type was largely captured by the eigengene values for Modules 1 and 2 (Fig. 2D,H and Supplemental Fig. S9A). The uninfected and *w*Mel-infected states were most significantly distinguished between Modules 9 and 10 (Fig. 2A,B,T and Supplemental Fig. S9A).

**Figure 3.**
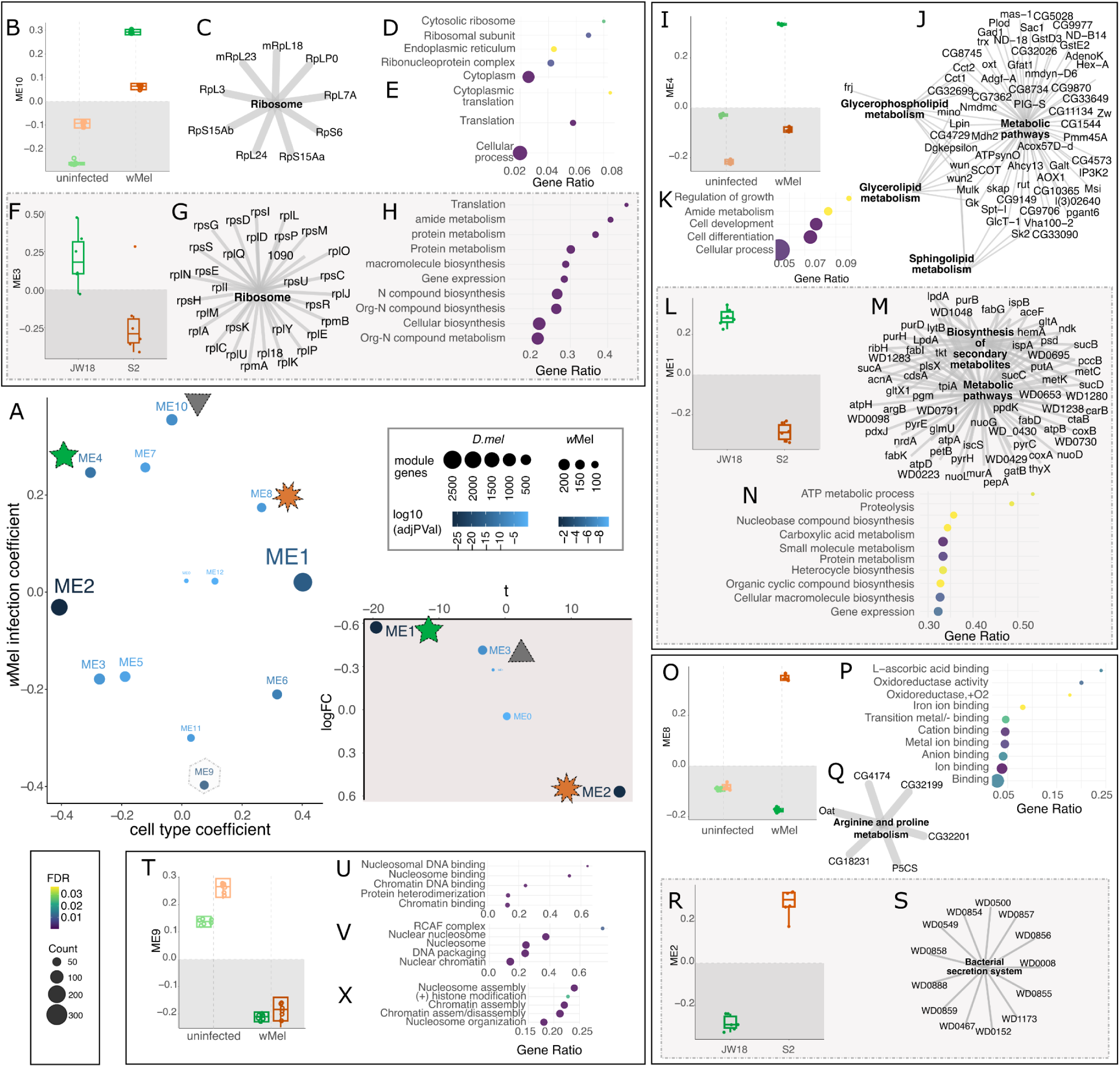
*Drosophila* and *Wolbachia* transcriptomes are functionally correlated. A) Coefficient estimates from linear models fit to (left) *D. melanogaster* and (right) *w*Mel WGCNA module eigengenes relative to their respective reference conditions. Left plot: The x– and y-axes contain the estimated linear model coefficients for the variables cell type and infection, respectively, for each module. Right plot: LogFC plots the direction and magnitude of eigengene expression. The t variable is the moderated t-statistic for the logFC difference between the two cell types for *w*Mel gene expression. Colored symbols indicate the correlated modules between transcriptomes: green star = JW18*w*Mel, orange star = S2*w*Mel, and grey triangle = *w*Mel-infection. The grey hexahedron outline = uninfected cells. B-S) Module eigengene evidence of cell type-specific functional correlation between *D. melanogaster* and *w*Mel transcriptomes. B-E,I-K,O-Q) *D. melanogaster* modules that correspond to F-H,L-N,R-S) wMel modules labeled in (A). T-X) Module 9 evidence associated with uninfected cells. Eigengene value plots for B,I,O,T) *D. melanogaster* and F,L,R) *w*Mel WGCNA modules. C,G,J,M,Q) KEGG enrichment network plots for each module (FDR-adjusted p-value<0.05). Gene ontology (GO) D,V) component, P,U) function, and E,H,K,N,X) process enrichment plots for each module, plotting the top 5-10 terms by gene ratio (FDR-adjusted p-value<0.05). Key: FDR p-value gradient and gene count dot size for GO plots.

The WGCNA modules most strongly associated with cell type and uninfected cell states were better annotated than the modules associated with infected cells, suggesting that *Wolbachia* infection may induce the expression of non-canonical host pathways (Fig. 3 and Supplemental Figs. S9-12). Modules 1, 2, 3, 5, 6, 9, and 11 were either solely associated with cell type or uninfected cells, or a subset of uninfected cells. Modules 4, 7, 8, and 10 were either associated with infected cells or a subset of infected cells. Despite both categories of modules containing a similar number of genes (173-2687 versus 240-556 genes, p=0.41 Wilcoxon Rank Sum Test), the uninfected modules were enriched in more known GO and KEGG terms than the modules associated with infected cells (120-1244 vs 14-68 terms, p=0.004, Wilcoxon Rank Sum Test; Supplemental Fig. S9 and Table 15). Modules 2-5, 7, and 9-12 contain genes that are in linkage along the genome (Supplemental Fig. S13), suggesting their transcription may be co-regulated.

### Transcriptomic analyses reveal *Wolbachia* genes correlated with host cell type

The *w*Mel strain of *Wolbachia* modulates its gene expression in response to the *in vitro D. melanogaster* cell line it infects (Fig. 3A and Supplemental Fig. S14A-C and Table S16). After quality control and processing, we detected 1,122 expressed *Wolbachia* genes among samples. Cell type corresponded to 128 genes with greater than 0.5x log2-fold change across replicates, with two having greater than 2x log2-fold change (p ≤ 0.01 FDR-corrected Wald Test). *Wolbachia* upregulated more genes in the S2 cell type than the JW18 cell type (108 vs 20 genes and 4 vs 0 genes at 0.5x and 2x log2-fold change, respectively).

GSEA analysis of the differentially expressed *w*Mel genes by model condition revealed nine bacterial functional pathways differentially regulated due to host cell type (p ≤ 0.05 permutation test; Supplemental Fig. S14D and Table S17). All of these pathways were enriched in S2 cells, consistent with S2 cells eliciting a larger differential expression response from *w*Mel than JW18 cells. Several of these *w*Mel pathways overlapped with differentially expressed host pathways (Supplemental Fig. S6I-J), including tRNA biosynthesis, glycolysis and other types of carbon metabolism, metabolite synthesis via oxocarboxylic acid metabolism and secondary metabolites, and ribosome biosynthesis. However, this approach failed to annotate the *w*Mel pathways upregulated in JW18 cells.

We inferred five signed WGCNA co-expression modules from the *w*Mel transcriptomes, three of which were significantly associated with the two *D. melanogaster* cell type conditions (p ≤ 0.05, Benjamini–Hochberg FDR-corrected moderated t-test; Fig. 3A, F-H, L-N, R-S, and Supplemental Figs. S14 and Tables S18-19). All three modules contained approximately one to two hundred genes each, which were significantly enriched in one or more KEGG pathways (p ≤ 0.05 Benjamini-Hochberg FDR-corrected hypergeometric test, Fig. 3G,M,S, Supplemental Fig. S15-16 and Table S20). Only 22-49% of genes in the three significant modules were differentially expressed (Fig. S14G), potentially due to natural disparities between cytoplasmic bacterial and eukaryotic mRNA abundances (Johnston and Bullman 2022; Imdahl et al. 2020). The two *w*Mel modules not associated with host cell type, Modules 0 and 4 were associated with ABC transporters and other membrane-associated proteins (Supplemental Fig. S15). The genomic locations of the genes in the modules were broadly distributed, with some clustering (Fig. S14H).

### Complementation between host and *w*Mel gene expression modules

The predicted functions of the three significant *w*Mel modules were correlated with the functions of co-occurring *D. melanogaster* modules (Fig. 3). The association between *w*Mel Modules 1 and 3, encoding ribosomal and metabolic pathways, respectively, and JW18 cells correlated with the functions of *D. melanogaster* Modules 10 and 4, and their association with the infected JW18 cell state (Fig. 3B-N). Functional correlation between *w*Mel and *D. melanogaster* modules in the S2 cell line condition was less explicit, as their annotations did not directly overlap, like they did for the JW18 cell line condition. However, comparing the S2-associated *w*Mel Module 2, which encodes a type IV secretion system, with the *D. melanogaster* modules associated with the infected S2 cell state suggest that the host must compensate for infection to a greater degree in this cell line (Fig. 3O-S and Supplemental Fig. S11A,C).

Translation machinery is upregulated in *w*Mel-infected JW18 cells, as well as in *w*Mel themselves when they infect JW18 cells (Fig. 3B-H), suggesting microbe and host interact at the translational level. Both JW18 and S2 cells were enriched in genes belonging to the ribosome and other translational machinery in the *w*Mel-infected state, relative to the uninfected state (Module 10 KEGG and GO enrichment p ≤ 0.05, Benjamini-Hochberg FDR-corrected hypergeometric test; Fig. 3C-E). Similarly, the *w*Mel transcriptome upregulated the expression of ribosomal, transfer RNA (tRNA), and other genes related to translation. (Module 3 KEGG and GO enrichment p ≤ 0.05, Benjamini-Hochberg FDR-corrected hypergeometric test; Fig. 3G-J). Finding *w*Mel infection correlated with altered translation is consistent with a previous high-throughput RNAi screen using these cells (Grobler et al. 2018).

Metabolic pathway expression is also paired between *D. melanogaster* and *w*Mel in JW18 cells (Figure 3I-N). The *w*Mel-infected JW18 cells expressed genes for growth and lipid metabolism (Module 4 KEGG and GO enrichment p ≤ 0.05, Benjamini-Hochberg FDR-corrected hypergeometric test; Fig. 3K-J) and repressed the expression of genes for microtubule-based transport for spindle assembly, chromatid segregation, and transcription factor complexes (Module 6 GO enrichment, p ≤ 0.05, Benjamini-Hochberg FDR-corrected hypergeometric test; Supplemental Fig. S11E-F). Altered glycerophospholipid metabolism suggests a mechanism underlying *w*Mel-induced changes in host cell morphology (Figure 1C-V), as this is the main constituent of plasma membranes (Lamari et al. 2025), and *w*Mel lacks a full pathway for glycerophospholipid synthesis (Molloy et al. 2016; Wu et al. 2004). Paired metabolic responses include pathways such as riboflavin synthesis (Supplemental Fig. S14C), which lead to nutritional mutualism in some contexts and strains (Ju et al. 2020; Hosokawa et al. 2010; Wiles et al. 2025; Moriyama et al. 2015). B-vitamin supplementation may explain how *w*Mel mitigates the costs of its presence in host cells, as insects cannot make these essential co-factors (Douglas 2017).

The transcriptional modules associated with S2 cells suggest that *w*Mel gene expression is costly in this host background and the *D. melanogaster* transcriptome mounts a counterresponse (Figure 3O-S and Supplemental Fig. S17). In addition to expressing genes for a type IV secretion system, the S2-associated wMel transcriptome was enriched in binary fission genes, including DNA translocase FtsK and septal ring protein RlpA (Supplemental Fig. S14A and Tables S15, S18). The S2 cell line upregulated arginine and proline metabolism in response to wMel infection (Fig. 4I, Supplemental Table S15, S18), suggesting the host responds to increased bacterial binary fission by synthesizing *w*Mel’s essential amino acids (Jiménez et al. 2019; Wu et al. 2004). Consistent with the hypothesis that *w*Mel compensates for lysis in S2 cells by upregulating binary fission, S2 cells rapidly clear many of the first *w*Mel to enter in *de novo* experimental infections (Supplemental Fig. S17) and *w*Mel titers are similar between stably-infected S2 and JW18 cells (Mirchandani et al. 2024). Capsid and subtilisin gene expression is also upregulated in S2 cells, and may reflect a pathogenic response from *w*Mel or it may reflect a de-repressed, stress response. Other upregulated *w*Mel genes in S2 cells include AAA-ATPases and conjugal transfer TrbC/type IV secretion proteins (Fig. 4B and Supplemental Tables S15, S18), which may be used to secrete effectors, such as the upregulated ankyrin repeat domain proteins, into the host cytoplasm. These activities are likely pathogenic to S2 cells (Jaboulay et al. 2021).

**Figure 4.**
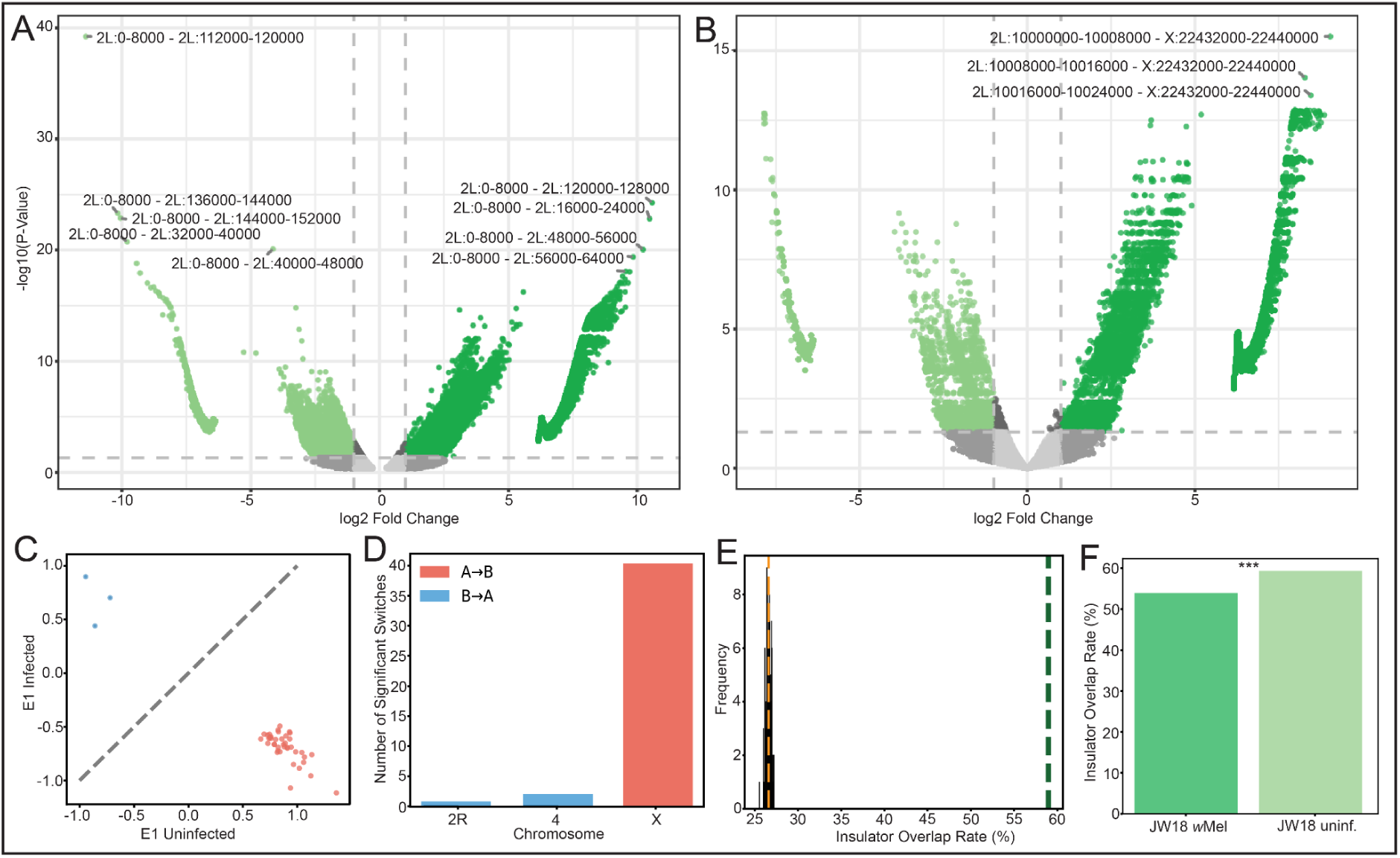
*Wolbachia* infection alters *Drosophila* chromatin contacts with X chromosome-specific compartmental switching. A,B) Volcano plots of differential A) cis and B) trans chromatin interactions identified by diffHic between *w*Mel-infected and uninfected JW18 cells. Positive log₂FC, stronger in *w*Mel; negative, stronger in uninfected. Dashed lines, significance thresholds (|log₂FC| > 1, FDR < 0.05). C) E1 eigenvector values for 1,851 50-kb bins in uninfected (x) vs *w*Mel-infected (y) cells. D) Bar plot of compartment switch counts per chromosome: 40/43 switches were A→B on the X chromosome. Significant switches (FDR < 0.1) colored red (A→B) or blue (B→A). E) Permutation null distribution (gray, n = 1,000) for insulator-differential contact overlap with observed (green) and expected (orange) rates indicated. F) Insulator positive overlap rates for differential contacts. Dark green, *w*Mel; light green, uninfected. ***p<0.001.

Overall, these results suggest that host cell transcriptomic state may influence the nature of *w*Mel gene expression, creating a feedback loop that shifts the interaction between *w*Mel and its host on a cell type-specific basis. The neuron-like nature of the JW18 cell line may present a hospitable environment for *w*Mel and may promote a more mutualistic-type of infection than seen in S2 cells. Consistent with their neuroblast-derived cytological properties (Grobler et al. 2018), JW18 cells express neuronal pathways such as inositol phosphate metabolism for intracellular signaling and calcium regulation (Acharya et al. 1998; Seeds et al. 2015) (Fig. 2G-P). This cell line does not appear to target *w*Mel for degradation, permitting *w*Mel to differentially express a suite of genes that may promote its intracellular persistence. In contrast, S2 cells express endocytosis and vesicle coat genes (Fig. 2C-G, Supplemental Fig. S14B, and Tables S3 and S13), suggesting they behave like hematopoietic-derived macrophages (Luhur et al. 2020; Krejčová et al. 2019; Kirkpatrick et al. 1995; Elwell and Engel 2005), and may be actively degrading *w*Mel *in vitro*. This macrophage-like nature may trigger a pathogenic transcriptional response from *Wolbachia* that subsequently induces a compensatory host response (Supplemental Fig. S11A-C,G-I).

The absence of *w*Mel infection is associated with the expression of host genes involved in histone and nucleosome modifications, suggesting that infection alters chromatin compaction (Figure 3T-X and Supplemental Fig. S10D-F and S11D-F). Both JW18 and S2 uninfected *D. melanogaster* cells upregulate chromatin, nucleosome, and histone modifiers (Fig. 3T-X), suggesting that infection alters chromatin compaction. Histone and chromatin modifiers are further upregulated in uninfected JW18 cells (Supplemental Fig. S10E-F), and repressed in the *w*Mel-infected condition (Supplemental Fig. S11E-F), consistent with the infection inducing larger phenotypic effects on the JW18 cell line than the S2 line (Fig. 1A-L).

### Infection is associated with altered chromatin contacts in JW18 cells

To investigate how *Wolbachia* infection affects host chromatin organization, we performed Micro-C chromatin conformation capture sequencing on JW18 *D. melanogaster* cells uninfected and infected with *w*Mel. We identified 262,206 significant differential interactions (FDR < 0.05), including 213,203 *cis* and 49,003 *trans* interactions (Fig. 4A-B, Supplemental Fig. S18, and 10.5061/dryad.ghx3ffc1g). As some genes had multiple differential contacts, the total count by gene was also informative: 9808 genes experienced differential chromatin contacts due to infection. Of these, 3064 genes exclusively gained contacts in the infected state, 22 genes exclusively lost contacts, and 6722 genes showed contact changes in both directions, suggesting widespread chromatin reorganization rather than uniform gain or loss. In cis, infected cells had 24,930 negative contact loop changes (less contact) relative to uninfected cells, and 188,273 positive (more contact). All of these cis changes occurred along the second chromosome, with nearly all (193,242) occurring along chromosomal arm 2L, consistent with the co-localization pattern of JW18-associated *D. melanogaster* WGCNA Module 11 genes (Supplemental Fig. S13). In trans, infection with *w*Mel altered 4,515 negative and 44,488 positive contact loops. All of these trans interactions involved chromosomal arm 2L, with 26,777 occurring with 3R (24,402 positive and 2,375.00 negative). Differential contacts with chromosomal 2L arm occurred second-most frequently with 3L, making up 18,300 interactions (16,535 positive and 1,765 negative). With the X-chromosome, chromosomal arm 2L had 2,799 differential interactions (2,542 positive and 257 negative) and chromosome 4 had 1,122 differential interactions (1,007 positive and 115 negative).

Infection was associated with altered contact probability across all three autosomes and the X chromosome (Supplemental Fig. S19A-F). Over genomic distances less than 100 bp, uninfected cells tended to have higher chromatin contact probabilities than infected cells (Supplemental Fig. S19G-L). While infected cells showed a visual trend toward reduced X chromosome contact density relative to uninfected cells, normalizing by autosomal contacts revealed this to be a modest 4.4% decrease, which was not statistically significant with the current sample size (Supplemental Fig. S20, mean ratio: infected = 0.448 ± 0.012, uninfected = 0.469 ± 0.008; Mann-Whitney U test, p = 0.67, n=2 per condition). However, given that the JW18 cells are male (Grobler et al. 2018) and the wMel strain encodes factors that interact with dosage compensation to facilitate male killing (Perlmutter et al. 2019), we explored chromatin contact alterations at other scales to determine whether infection impacts dosage compensation *in vitro*.

### X-linked chromatin compartments switch to the inactive state and TADs are lost during wMel infection

The chromosome compartments that switch during wMel infection are largely restricted to the X chromosome and predominantly involve chromatin compaction (Fig. 4C-D, Supplemental Fig. S21 and Table S21). Eukaryotic genomes are organized into transcriptionally active A compartments and transcriptionally silent B compartments (Hildebrand and Dekker 2020). Analysis of genome A/B compartment identity across 1,851 genomic bins showed reorganization during *w*Mel infection, with 22 compartments (1.2%) switching between conditions. Notably, 21 of these 22 changes (95%) were A to B transitions on the X chromosome, while only three B to A switches occurred on autosomes.

We quantified the changes due to infection at topologically-associating domain (TAD) boundaries, the chromatin loops within TADs, and hotspots of conformation change along the *D. melanogaster* chromosomes, revealing a widespread pattern of decreased regulation (Supplemental Fig. S22). TADs represent sub-megabase domains of preferential chromatin contact that correlate with regulatory landscapes (Dixon et al. 2012; Ramírez et al. 2018). We identified interacting regions of the genome by calculating sliding-diamond window insulation scores along the contact maps (Supplemental Table S22). From these, we identified windows with the largest difference between the minimum and maximum insulation score as boundary regions between TADs. Total TAD boundaries decreased with *w*Mel infection, from 161 TAD boundaries in uninfected cells to 136 boundaries in infected cells (Supplemental Fig. S22C). Across all chromosomes, uninfected and infected cells shared 124 TADs (72%), with 37 unique to the uninfected state and 12 unique to wMel-infected cells (Supplemental Fig. S22D and Table S23). While these TADs were differentially detected between conditions, only one TAD from 1.5 to 1.6 Mb on 3L significantly switched boundary states (p=0.0099, two-sided z-score of boundary strength differences, centred at 0). Interestingly, this region overlapped the start of 3L that is enriched in infection-associated Module 10 genes (Supplemental Fig. S13 and Fig. 3A,B). Within TADs, we identified 203 loop changes associated with infection (q < 0.01, cooltools dot-calling), of which 147 were lost loops and 56 were gained loops (Supplemental Fig. S23A,D). As suggested by altered TAD boundaries and loops, hotspot analysis revealed differential contacts to be non-randomly distributed across the genome, with 31 regions exhibiting significant reductions in contact density due to infection (p = 4.693e⁻-13, permutation test, Supplemental Fig. S23B-D).

### Differential chromatin contacts are enriched at insulator protein binding sites

We tested for regulatory element term enrichment in our differential contact data to better understand the upstream and downstream implications of increased compartment compaction and decreased TAD/loop structural regulation in JW18 cells due to *w*Mel infection, finding enrichment at insulator protein binding sites (Fig. 4E-F and 10.5061/dryad.ghx3ffc1g). Insulator-binding factors such as CTCF, dCTCF, Su(Hw), and BEAF-32, establish and maintain chromatin domain boundaries in *Drosophila* (Ulianov et al. 2021). Comparing the overlap of differential contacts with annotated insulator protein binding sites revealed a significant enrichment compared to random permutations (p<0.001, Fig. 4E). Uninfected cells were enriched in differential positive contacts at these sites, compared to *w*Mel-infected cells (p=2.16e-15, MannWhitney U-test, Fig. 4F). While enhancer-enhancer and enhancer-promoter contact frequencies were significantly enriched in infected compared to uninfected cells, their logFC and effect sizes were below significance thresholds (Supplemental Fig. S24). In total, these results suggest that infection-mediated changes in chromatin contacts occur preferentially at sites occupied by architectural insulator proteins, opposed to enhancers, leading to changes in loop and TAD organization genome-wide.

The abundance of X-linked A-to-B compartment switches induced by *w*Mel infection and the decrease in insulator-contact overlap rate, leading to loop and TAD loss, suggests that this strain may interact with the dosage compensation apparatus *in vitro,* similar to other *Wolbachia* strains (Harumoto et al. 2018; Arai et al. 2025; Lefoulon et al. 2025). Dosage compensation is achieved in *Drosophila* through upregulation of the single male X to match the expression from two X chromosomes in females (Xie et al. 2025). This process is mediated by the Male-Specific Lethal (MSL) complex, and involves histone H4 lysine 16 acetylation (H4K16ac), which leads to a more open X chromosome chromatin state (Harumoto et al. 2018; Straub et al. 2005; Pal et al. 2023). Transgenic expression of wMel’s *WO-mediated killing (wmk)* gene in absence of *w*Mel infection is sufficient to induce dysregulation of the male dosage compensation apparatus and DNA damage to the X chromosome in embryos (Perlmutter et al. 2019). While male killing does not naturally occur in *w*Mel infections, the *w*Bif strain from *Drosophila bifasciata* accomplishes male killing via targeting the dosage-compensated chromosome (Harumoto et al. 2018). Therefore, we were motivated to look for further evidence that this largely X-linked chromatin phenotype may be related to dosage compensation manipulation *in vitro*.

### Increased chromatin interactions at the *ndf* locus and H3K36me-marked regions in *w*Mel-infected cells

The most significant trans differential contact we detected between infected and uninfected cells was between the *nucleosome-destabilizing factor* (*ndf*) gene on the second chromosome and several regions along the X chromosome (FDR, q=2.94E-12, Fig 4B), which provided a mechanistically testable hypothesis for *w*Mel-mediated alterations in the dosage compensation pathway. Ndf functions as a recruitment factor for the MSL Dosage Compensation Complex (DCC), physically connecting histone H3 trimethylated at lysine 36-(H3K36me3-) marked chromatin to MSL proteins (Fei et al. 2022, 2018) (illustrated in Fig. 5A). The Ndf protein is a chromatin reader containing a PWWP domain that specifically recognizes H3K36me3. This mark is cotranscriptionally deposited along gene bodies genome-wide by Set2 histone methyltransferase (Bell et al. 2008; Walshe et al. 2025). Following initial recruitment of the MSL complex to the X chromosome by co-transcriptional assembly at *roX* genes and subsequent recognition of MSL recognition elements (MREs) at high-affinity sites/chromatin entry sites (HAS/CES), Ndf facilitates spreading to H3K36me3-marked active genes, where the DCC complex deposits H4K16ac to upregulate transcription approximately two-fold (Gelbart and Kuroda 2009).

**Figure 5.**
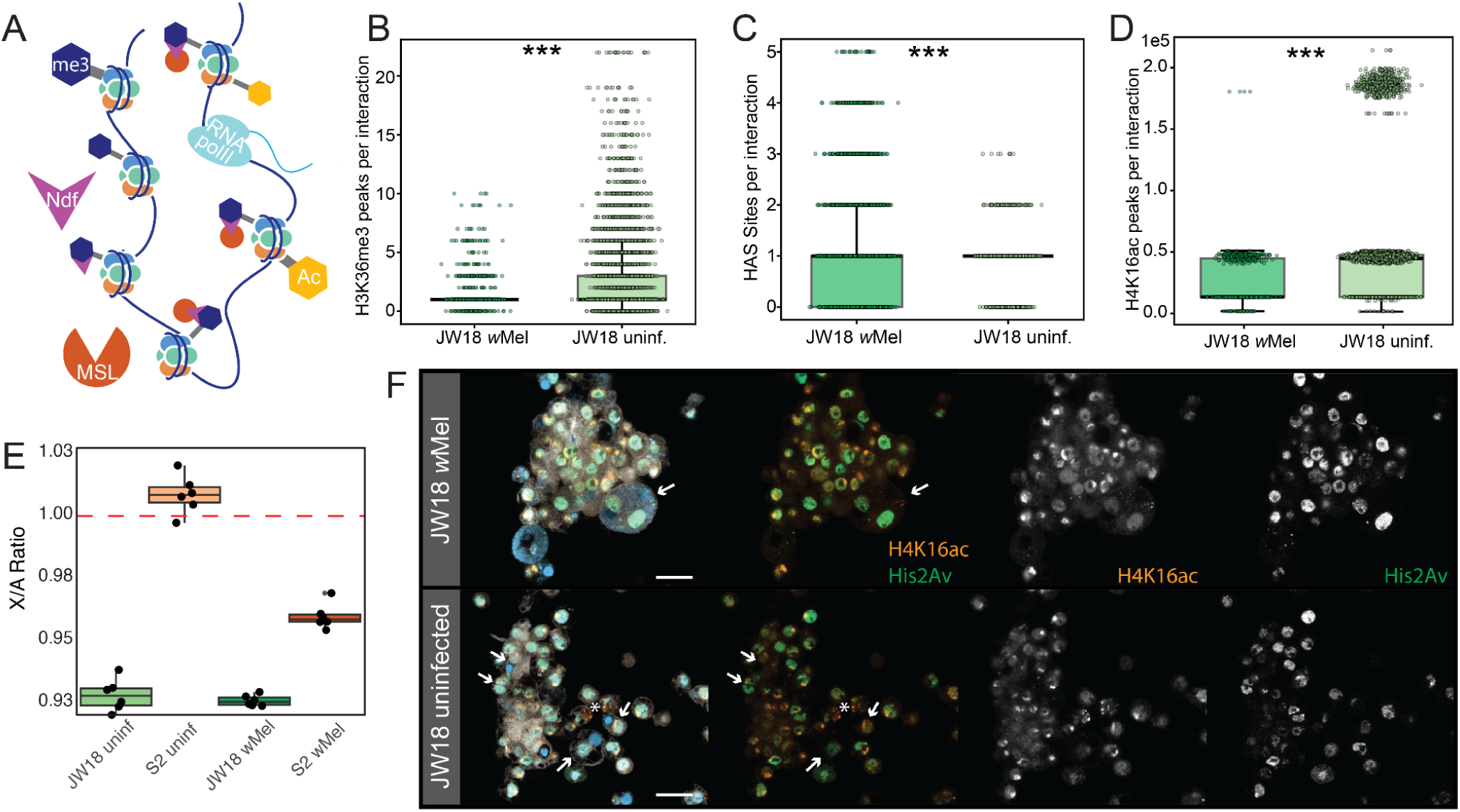
*w*Mel infection alters X chromosome contacts related to dosage compensation, without downstream impacts. A) Schematic of MSL-mediated dosage compensation. Ndf binds H3K36me3 (me3) on active gene bodies and recruits MSL, which deposits H4K16ac (Ac) via MOF, opening chromatin for RNA Pol II upregulation. Significant X-chromsome positive differential contact-interaction rates for B) H3K36me3 ChIP-seq peaks C) HAS/CES sites, and D) H4K16ac peaks on the X chromosome. E) X-to-autosome transcriptional ratio. Error bars, SEM. F) Confocal images of immunolabeled JW18 cells (top) infected and (bottom) uninfected with *w*Mel. Panel columns reflect fluorescent channels: Far left panel: merged Jupiter-GFP (grey), DAPI (skyblue), H4K16ac (vermillion), and pHis2Av (bluish-green). Other panels are the channels indicated in the overlay.

Differential contacts were enriched at H3K36me3 peaks along the X chromosome (p<2.2e-308 permutation test, Fig. 5B), with a lower H3K36me3-positive contact overlap rate in infected cells than uninfected cells (81.0% versus 87.9%%, p=3.15e-06, one-sided binomial test, Supplemental Fig. S25A), suggesting *w*Mel interferes with this chromatin regulatory network. While Ndf has functions elsewhere in the genome to destabilize nucleosomes in an ATP-independent manner and facilitate RNA polymerase II transcription (Fei et al. 2018), it is not yet clear if there is a mechanism enabling early and biased Ndf-localization to the X chromosome during *Drosophila* development. Our results suggest that physical proximity between the *ndf* locus on 2L and X-linked H3K36me3 sites may enable such a mechanism, which is disrupted in infected cells.

### Altered chromatin contacts at dosage compensation complex chromatin entry sites

Differential chromatin contacts were enriched at DCC entry sites along the X chromosome (p<2.2e-308 permutation test, Fig. 5C), with a higher HAS/CES-positive contact overlap rate in infected cells than uninfected cells (81.9% versus 74.4%, p=2.91e-7 one-sided binomial test, Supplemental Fig. S25B), further suggesting that wMel alters host dosage-compensation pathways at the chromatin level. Before interacting with transcriptionally placed H3K36me3 marks, the DCC first spreads across the 150-250 HAS/CESs encoded along the single male X chromosome, which are known to closely physically associate in male chromatin (Grimaud and Becker 2009), enabling spread from one HAS/CES to the next. These sites are characterized by a specific DNA sequence motif termed a MSL recognition element (MRE), and are preferentially located within or near H3K36me3-marked gene bodies (Alekseyenko et al. 2008). In male *Drosophila* undergoing normal dosage compensation, HAS/CES act as nucleation centers from which the MSL complex spreads in cis to nearby active genes to ensure proper dosage compensation across the X chromosome (Straub et al. 2005; Wang et al. 2013). The enrichment for increased contact at HAS/CES sites is consistent with infection altering processes involved in initiating dosage compensation at the chromatin-level.

### Reduced local chromatin contacts in H4K16ac-enriched regions

Differential chromatin contacts were enriched at acetylated H4K16 sites along the X chromosome (p<2.2e-308 permutation test, Supplemental Fig. S25C), with fewer H4K16ac peaks per interaction in infected cells than uninfected cells (mean 2412.5 versus 4365.4, p=3.75e-57 Mann-Whitney U, Fig. 5D).The MSL complex contains Males absent On the First (MOF), a histone acetyltransferase that marks histone H4 at lysine 16 (H4K16ac) (Wei et al. 2022; Straub et al. 2005). H4K16ac is the key effector modification of dosage compensation through disruption of higher-order chromatin compaction by preventing inter-nucleosomal interactions and recruits chromatin remodeling factors that upregulate X-linked transcription (Harumoto et al. 2018; Gelbart and Kuroda 2009). Overall, finding elevated chromatin contacts at the sites of acetylation that initiate male X-chromosome decompaction, with fewer positive contacts in infected cells than uninfected, suggests that wMel infection may weaken dosage compensation downstream of HAS/CES entry.

### Doubly-compensated X-linked dosage

The changes in chromatin contacts suggestive of disrupted dosage compensation in JW18 cells (Fig. 4,5A-D) did not persist at the transcriptional level (Fig 5E-F). Infection increased chromatin contacts at CES/HAS sites along the X (Fig. 5C), between 2L-linked *ndf* and positions along the X (Fig. 4B), and increased the proportion of inactive compartments (Fig. 4C-D), while reducing the proportion of positive contact at H3K36me3 and H4K16ac sites (Fig. 5B,D). These data suggest that *w*Mel interacts with host dosage compensation mechanisms, but repressive and depressive pathways may cancel each other out. To assess the downstream impacts of *w*Mel infection on dosage compensation we quantified the *D. melanogaster* X-to-autosome ratio of transcript expression from the bulk RNAseq data by cell line and infection, finding that the cells all exhibited an X:A ratio close to 1.0 (0.93-1.0, Fig. 5E). While the S2 cell X:A ratio did decline upon infection (from 1.01 to 0.96, q=1.157e-06 BH-corrected Welch’s t-test), the JW18 ratio was consistently 0.93.

At the protein level, both uninfected and infected JW18 cells exhibit nuclear anti-H4K16ac staining, with strong localization to a segment of the genome consistent with the X chromosome (Fig. 5F, and see (Aleman et al. 2021)). There may be differences in the timing of H4K16ac deposition after mitosis, as the uninfected cells we observed in mitosis exhibited the X-specific H4K16ac staining pattern before telophase was complete, whereas infected cells were did not express this factor until after this stage (arrows in Fig. 5F). Uninfected cells also exhibited asymmetric His2Av staining patterns between daughter cells at this stage of mitosis, whereas infected cells appeared more symmetrical. Known *Wolbachia* mechanisms for male-killing via hijacked dosage compensation involve DNA damage to the compensated chromosome (Harumoto et al. 2018), therefore we assessed the staining pattern of phosphorylated Histone H2A gamma variant (pHis2Av), which is phosphorylated due to DNA damage (Lake et al. 2013). Both uninfected and infected cells had a general nuclear pattern of pHis2Av localization, without apparent bias for the H2K16ac-labeled X-chromosome (Fig. 5F), indicating that downstream dosage compensation is unperturbed.

## Discussion

This study demonstrates that the relationship between *Wolbachia* and its *Drosophila* host is deeply cell-type-specific, operating through transcriptional and chromatin-level mechanisms that are hard-coded into the epigenetic landscape of the host cell. Using matched infected and uninfected somatic cell lines as a tractable *in vitro* system, we show that *w*Mel does not impose a uniform response to infection in its host. Instead, *w*Mel responds to the identity of the cell it occupies, inducing complementary changes in both bacterial and eukaryotic gene expression that appear to shift the fundamental cost-benefit dynamics of infection. These findings may explain how *Wolbachia* infectivity is altered by the host tissue type they were isolated from (Pigeault et al. 2026).

The divergent phenotypic and transcriptomic responses of JW18 and S2 cells to *w*Mel infection illustrate the possible spectrum of cell-type-specific responses to infection. In the neuroblast-like JW18 background, *w*Mel upregulates ribosomal machinery and metabolic pathways, including riboflavin synthesis, in concert with a reciprocal host transcriptional response, consistent with mutualistic interactions in which the bacterium supplements host nutritional requirements and supports cellular persistence (Douglas 2017; Lindsey et al. 2025; Hosokawa et al. 2010). In the macrophage-like S2 background, the picture reverses: *w*Mel upregulates binary fission and secretion machinery, while the host mounts a compensatory metabolic response, suggesting active bacterial replication under immune pressure followed by partial clearance. These contrasting profiles argue that the same bacterial strain can adopt fundamentally different interaction modes depending solely on the molecular environment provided by the host cell. This finding has direct implications for interpreting the context-dependent phenotypes that make *Wolbachia* both a biological control tool and an evolutionarily plastic symbiont.

Chromatin contact sequencing corroborates our transcriptomic evidence that *w*Mel infections alter host epigenetics, and corroborates prior evidence that *w*Mel encodes proteins for interacting with host chromatin (Terretaz et al. 2023) and interfering with host dosage compensation (Perlmutter et al. 2019). The near-exclusive A-to-B compartment switching on the X chromosome, combined with altered contacts at HAS/CES dosage compensation entry sites, the *ndf* locus, H3K36me3-marked gene bodies, and H4K16ac sites, provides the first evidence that *w*Mel engages with the dosage compensation machinery at the chromatin level *in vitro*. That these upstream perturbations do not propagate to a detectable change in the X-to-autosome transcriptional ratio suggests that repressive and activating chromatin forces may be balanced through a form of epigenetic buffering, rather than silencing. This nuanced result aligns with the observation that *w*Mel does not naturally induce male killing in *D. melanogaster*, even though the *wmk* gene it encodes is sufficient to dysregulate dosage compensation when expressed transgenically (Perlmutter et al. 2019). The *in vitro* chromatin phenotype we describe may represent a latent, buffered interaction with dosage compensation pathways that is only realized as a lethal phenotype in specific genetic or developmental contexts.

The cell-type specificity of *w*Mel’s transcriptional, phenotypic, and chromatin effects suggests that the diverse phenotypes induced by *Wolbachia* across host tissues and species may emerge from the intersection of each *Wolbachia* strain’s genetic repertoire with distinct host cell states. Dissecting these intersections systematically, offers a tractable path toward resolving the long-standing challenge of identifying the genetic and mechanistic basis for *Wolbachia*-induced host phenotypes.

## Methods

### Materials and Methods

#### Wolbachia infected Drosophila cell culture

We created paired wMel-infected and uninfected *Drosophila melanogaster* S2 (Schneider 1972) and JW18 (Serbus et al. 2012b) following (Lum et al. 2026), and confirmed infection status using fluorescence in situ hybridization (FISH) with DNA oligonucleotide probes targeting the *Wolbachia* 16S ribosomal RNA (as in (White et al. 2017)). We quantified the relative *w*Mel:*D. melanogaster* genomic titer with Illumina shotgun sequencing (Mirchandani et al. 2024). We quantified cell line doubling rates when splitting and seeding cells into new flasks. (Supplemental Materials, Sections 1-3 and (Mirchandani et al. 2024)).

#### Live phenotypic imaging and analysis

We imaged 96-well plates of wMel-infected and uninfected JW18 and S2 cells stained with NucBlue Live ReadyProbes Reagent (Invitrogen, R37605) on an Opera Phenix High-Content Screening System (PerkinElmer/Revvity) with a 20X objective. We used Harmony software to perform cell segmentation and measure cell area, contact area, distance between cells, and fluorescence intensity, and tested for statistical differences in these measurements using a Welch’s two-tailed t-test (Supplemental Materials, Section 4 and Table 1).

#### Dual bulk transcriptomics

*D. melanogaster* cells stably infected with *Wolbachia* were harvested from confluent cultures at 23°C, pelted by centrifugation (16,000xg, 10 min, 4°C), and stored at –80°C. Following pellet lysis, RNA was extracted, ribosomal RNA depleted using sequential eukaryotic and bacterial rRNA depletion kits (Qiagen FastSelect), and cDNA synthesised using random hexamers. Dual-indexed Illumina libraries were prepared from these cDNAs and sequenced as 2×150bp reads on a NovaSeq (Supplemental Materials, Section 5a and Table 2).

We trimmed adapter sequences from demultiplexed reads using Trimmomatic (v0.39), and quantified transcripts by pseudoalignment with Kallisto (v0.45.1) against a merged host-symbiont reference transcriptome combining wMel (GCF_000008025.1) and *D. melanogaster* (GCF_000001215.4) assemblies, with simultaneous dual-genome mapping to prevent cross-species mismapping (Supplemental Materials, Section 5b, and Supplemental Table S24). Host and symbiont transcriptomes were separated prior to normalisation, and *D. melanogaster* alignments were imported into R with Tximport and processed in DESeq2 (Love et al. 2014) using a design accounting for cell type, infection status, and their interaction. We removed low-count genes by requiring a minimum read count of 10, 20, or 70 across all six samples per condition for differential expression, bulk-to-single cell clustering, or WGCNA analyses, respectively (Supplemental Materials, Section 5c).

We filtered, normalized, and clustered the bulk transcriptomic data with the *Drosophila* cell atlas using the Leiden algorithm in Scanpy (Wolf et al. 2018). We reduced dimensionality of the data by UMAP and PCA techniques, and identified marker genes with Wilcoxon rank-sum differential expression analysis. Bulk RNA-seq data were then integrated with three reference single-cell atlases (Fly Cell Atlas, Embryo Atlas, and Myeloid Blood Cell Atlas) using BBKNN batch correction on shared genes. We performed cell type classification using distance-weighted kNN and assessed confidence via permutation testing (1000 iterations). UMAP visualisations were generated to display bulk samples in the context of reference cell types across all three atlases (Supplemental Materials, Section 5d).

We tested for differential expression in DESeq2 using Wald tests with Benjamini-Hochberg correction, modelling host transcriptomes as a function of cell line, infection, and their interaction, and *Wolbachia* transcriptomes as a function of cell line alone. This yielded 10,839 expressed host and 1,122 expressed symbiont transcripts after low-count filtering (Supplemental Materials, Section 5e). We ranked differentially expressed genes output from DESeq2 by stat value and used for GSEA in clusterProfiler, including KEGG and GO enrichment analyses (Supplemental Materials, Section 5f). For WGCNA, we used filtered and rlog-normalised counts to construct signed co-expression networks, yielding 13 host and 5 symbiont eigengene modules whose associations with experimental conditions were tested by linear regression in limma. We subjected module gene lists to GO enrichment analysis using clusterProfiler, ShinyGO, and PANGEA (Supplemental Materials, Section 5g).

#### *Drosophila* cell atlas *Wolbachia* titer estimation

We assessed the *Wolbachia* infection status of the three *Drosophila* cell atlas datasets used for bulk-to-atlas mapping by aligning subsampled reads (1,000,000 reads per sample) to a combined *D. melanogaster*/*w*Mel reference genome using STAR (Dobin et al. 2013). We quantified gene expression by featureCounts (Liao et al. 2014). We exploited poly-T mispriming to rRNA regions (similar to (Robinson et al. 2024)) to estimate infection status by comparing the ratio of *w*Mel to *D. melanogaster* ribosomal RNA reads against known infected and uninfected scRNAseq controls (PRJNA788731) (Dou et al. 2021) using Fisher’s exact test, with samples classified as infected, uninfected, or inconclusive based on their metadata. We estimated 95% confidence intervals for *w*Mel/*D. melanogaster* rRNA ratios by bootstrap analysis (Supplemental Materials, Section 6).

#### Micro-C Chromatin Conformation Capture

We performed Micro-C chromatin capture on two biological replicates per condition using the Dovetail Genomics Micro-C kit, with dual crosslinking (DSG and formaldehyde), MNase digestion, and paired-end Illumina sequencing (NovaSeq S4, 2×150bp) (Supplemental Materials, Section 7a-b and Table 3). We aligned trimmed reads to the *D. melanogaster* reference genome with BWA-MEM (Li 2013) and processed the alignments with pairtools to identify ligation junctions and remove duplicates. Chromatin contact maps were constructed at 1kb resolution using cooler (v 0.9.3) (Abdennur and Mirny 2019) (Supplemental Materials, Section 7c). Differential chromatin interactions were identified with diffHic following TMM normalisation, using a likelihood ratio test for infection status at FDR < 0.05 (Supplemental Materials, Section 7d). To prevent structural variants (SVs) in the JW18 cell line from confounding contact analysis, we performed Nanopore long-read sequencing and identified SVs with SVIM (Supplemental Materials, Section 7e). Differential chromatin interactions overlapping SV breakpoints were excluded from downstream analyses.

We performed enrichment analyses to test for significant changes in chromatin topology and associations between differential contacts and annotated elements in the *D. melanogaster* genome (Supplemental Materials, Section 7f-h). A/B compartment analysis was performed at 50kb resolution using cooltools eigenvector decomposition, with compartment switches assessed against a permutation null model. TAD boundaries and chromatin loops were identified via insulation score profiling and cooltools.dots, and differential-interaction hotspots were mapped by aggregating significant interactions into 50kb bins. The enrichment analyses we performed were motivated by the number of interactions we detected that involved the X chromosome. Enrichment analyses tested whether differential interactions were overrepresented at insulator binding sites (Class I/II, from modENCODE ChIP-seq), enhancer regions (housekeeping and developmental), histone modification peaks (H3K36me3, H4K16ac), and dosage compensation high-affinity sites (HAS/CES), using combinations of binomial tests, permutation testing, Fisher’s exact test, and chi-square tests with Benjamini-Hochberg correction throughout.

#### Immunocytochemistry

We seeded cells on no. 1.5 glass coverslips in 6-well plates (Corning 3516) the day prior to staining. Cells were fixed in 8% paraformaldehyde/PBS for 12 min at RT, washed three times in PBS-T, blocked in 1% BSA/PBS-T for 1 hr, then incubated with primary antibodies for 1 hr at RT: anti-phHis2Av (UNC93-5.2.1, DSHB) at 1:500 and anti-H4K16ac (600-401-J00, Rockland) at 1:100. After three PBS-T washes, cells were incubated with secondary antibodies (1:500 in PBS-T) for 1 hr at RT and mounted in VECTASHIELD+DAPI (H-1200). Images were acquired on a Leica Stellaris confocal microscope with a 63x objective at Nyquist sampling and processed in ImageJ.

#### Data and code availability

All scripts are available from our Github pages: https://github.com/shelbirussell/Dual_bulk_RNAseq_analysis-Jacobs_et_al.git

https://github.com/jodiejacobs/wolbachia_induced_differentiation.git

Short and long read data can be found under NCBI BioProject PRJNA1240446.

Analysis results are on Dryad. https://doi.org/10.5061/dryad.ghx3ffc1g.

Reviewer link: http://datadryad.org/share/0FDVQailfgQ9rEaD6CE9mvFVoJ7x2yEmUzDCHY5s-ww

## Acknowledgements

We thank the UCSC Life Sciences Microscopy Center (RRID:SCR_021135) and Ben Abrams for training and the use of their microscopes. Phenix technical support was provided by Beverly Rabbits from the UCSC Chemical Screening Center RRID:SCR_021114. This work was supported by UC Santa Cruz and the NIH: R00GM135583 and R35GM157189 to SLR; T32HG012344 to JJ; and S10 Grants 1S10OD036314-01A1 to RRID:SCR_021135 for the Leica Stellaris and 1S10OD028730-01A1 to RRID:SCR_021114 for the Phenix Imager.

## Declaration of Interests

Richard E Green holds patents for the methods behind the Micro-C chromatin conformation capture kit and he holds a financial stake in the company who sells these kits, Dovetail Genomics. No other authors have any financial interests or patents to declare. None of the authors were board or advisory committee members or paid consultants related to this study or journal.

## Notes

### Summary of Updates

This version incorporates experiments and analyses motivated by comments from a first round of peer review.

