## Supplemental Methods for "*Wolbachia* infection induces host cell state changes *in vitro* and determines symbiotic fate in *Drosophila*"

### Supplemental Text Contents:

Experimental Methods ……….. p. 1 - 22
Resources Table ………………. p. 23 - 25
Supplemental References …… p. 26 - 30

### Experimental Methods

1. Drosophila cell culture

We created paired infected and uninfected cell lines at comparable split passages by curing *w*Mel-infected S2 (Schneider 1972) and JW18 (Serbus et al. 2012) with 10 mM (10 µg/mL) doxycycline for 10 weeks, followed by 10 weeks recovery from antibiotic treatment, following (Lum et al. 2026).

We maintained *Wolbachia w*Mel-infected and uninfected immortalized *Drosophila melanogaster* cell lines in 4 mL of Shields and Sang M3 Insect Medium (MilliporeSigma S3652) supplemented with 10% v/v Fetal Bovine Serum (FBS, ThermoFisher A3160502) in plug-seal T-25 flasks (Corning 430168) in a refrigerated incubator set to 23°C (Mirchandani et al. 2024b).

Cell cultures were maintained on a seven-day schedule, splitting at a 1:2 ratio for *w*Mel-infected lines and 1:6 to 1:4 ratio for uninfected lines. For passaging, we removed adherent cells by scraping the flask surface with a sterile bent Pasteur pipette (Fisher, 1367820D) after aspirating the media. Cells were then resuspended in fresh media, transferred at the appropriate ratio to new T25 flasks, and the volume adjusted to 4 mL with fresh media.

2. Validation of *Wolbachia* infections

To validate *Wolbachia* infections in our cell culture lines, we employed methods previously described in (Mirchandani et al. 2024b). Briefly, infections were confirmed using fluorescence in situ hybridization (FISH) with DNA oligonucleotide probes targeting the *Wolbachia* 16S ribosomal RNA. Cells were fixed in paraformaldehyde, subjected to a series of hybridization and washing steps, counterstained with DAPI, and imaged either mounted in Vectashield mounting media on a Leica Widefield epifluorescent microscope or imaged in 1xPBS in a 6-well dish on a Leica DMi8 epifluorescent microscope. For quantitative assessment of *Wolbachia* infection by genomic titer, we collected 1.5 mL of cell culture and performed Tn5 Illumina library preparation following protocols detailed in (Mirchandani et al. 2024b, 2024a).

3. <u>Cell growth rate experiments</u>

   We quantified cell line doubling rates when splitting and seeding cells into new flasks with a hemocytometer and Millicell Digital Cell Imager. While handling the cell lines as described above for "Cell Culture Maintenance", we retained a few dozen microliters for sampling cell concentration after pipetting the final split dilution into a fresh flask. We quantified these samples by counting cells in a 10 uL volume (X number of cells (>100) measured per Y number of boxes (>1 if <100 cells/box) * W dilution factor (2 if diluted by ½) * 10,000 mL-1 = Z number of cells/mL). This process was repeated one week later, except cells were resuspended by scraping prior to media removal so that the week's worth of growth could be quantified. Cells were then diluted as described above for normal maintenance. This was repeated every week for six weeks to obtain the data in Fig. 1B.

4. <u>Phenix imaging and Harmony analysis</u>

   We seeded *w*Mel-infected and uninfected JW18 and S2 cells into 96-well imaging plates (100 µL per well) in Shields and Sang M3 medium supplemented with 10% FBS. We stained *Wolbachia* and nuclear DNA with NucBlue Live ReadyProbes Reagent (Invitrogen, R37605; Hoechst 33342 ex/em 360/460 nm) at 2 drops per mL. We replaced the culture medium with the dye-containing medium for the incubation period, and then replaced it with fresh medium prior to imaging. Plates were imaged on an Opera Phenix High-Content Screening System (PerkinElmer/Revvity) using a 20X (120 ms exposure, 50% excitation power), acquiring brightfield, Hoechst, and GFP channels. Revvity's Harmony software performed cell segmentation, as well as measurements of cell area, contact area, distance between cells, and fluorescence intensity. We tested for statistical differences in these measurements between cell types and infection states with Welch's two-tailed t-test from stats.ttest_ind.

5. Dual bulk transcriptomics

a. *Sample collection and sequencing*

We collected immortalized *Drosophila melanogaster* cell culture cells stably infected with *Wolbachia* from confluent cultures grown at 23°C. For each sample, 1.2 mL (at ~2e6 cells/mL) of cells were pelleted by centrifugation at 16,000xg for 10 minutes at 4°C. Following supernatant removal, we promptly transferred the cell pellets to -80°C for storage.

Frozen cell pellets were shipped on dry ice to Genewiz Azenta Life Sciences for RNA extraction, cDNA synthesis, Illumina library preparation, and Illumina sequencing. The cell pellets were lysed, ribosomal RNA sequences were depleted with sequential eukaryotic and bacterial rRNA depletion kits (Qiagen FastSelect), and the remaining RNAs used as templates for cDNA synthesis using random hexamers. Illumina dual-indexed libraries were made from these cDNAs and sequenced as 2x150bp reads on a NovaSeq.

b. *Read processing and pseudoalignment*

We processed and analyzed RNAseq datasets for unsupervised clustering and differential expression analyses using standard computational approaches and custom parsing scripts. We trimmed adapter fragments from the demultiplexed RNAseq reads with Trimmomatic (v 0.39) (Bolger et al. 2014). We quantified transcripts by pseudoalignment with Kallisto (Bray et al. 2016). To obtain alignments against the full, non-redundant host-symbiont transcriptome, we merged the NCBI RefSeq assemblies for the *w*Mel reference genome CDSs and RNAs from genomic (accession GCF_000008025.1) and the *D. melanogaster* reference genome RNAs from genomic (accession GCF_000001215.4; Release_6_plus_ISO1_MT) as in (Russell et al. 2023). Simultaneous mapping to both genomes was performed to avoid cross-species mismapping (Chung et al. 2021). This reference transcriptome was indexed at a k-mer length of 31 in Kallisto (version 0.45.1) (Bray et al. 2016) and reads were pseudoaligned against this reference with the “kallisto quant” command and default parameters. Host and symbionts have distinct transcriptome distributions (Marsh et al. 2017), necessitating the separation of the two transcriptomes prior to transcript normalization and quantification in DESeq2 (Love et al. 2014), which we performed with a custom script.

c. *Transcriptome normalization and quantification*

We imported the subset *D. melanogaster* Kallisto transcriptome alignments into R with Tximport (Soneson et al. 2016). We estimated gene-level normalized counts by mapping transcript-level abundances to gene IDs (see Supplemental Table 24). Using DEseq2's DEseqDataSetFromTximport, we calculated the gene-level offset that corrects for average transcript length across samples with design = ~cell type + infection + cell type:infection. For each transcriptome, we filtered out low-count and low-coverage genes across samples by requiring all six samples of each cell type and infection state to have a minimum read count of 10, 20, or 70, for differential expression, bulk-to-single cell clustering, or weighted gene co-expression network analysis (WGCNA), respectively. This gene count matrix was exported as a .tsv file for bulk-to-single cell clustering.

d. *Bulk to Single Cell Clustering*

To analyze the effects of immortalization and *Wolbachia* infections on host cell identity, we performed two unsupervised clustering analyses of the pseudoaligned gene counts output from Kallisto. We imported the gene count matrix .tsv file from DEseq2 into Scanpy (Wolf et al. 2018) as an AnnData object. The sample by gene matrix was filtered to retain samples with at least 200 genes expressed and genes expressed in at least three samples. Counts based normalization was performed to a target sum of 1e4 gene counts per sample.

We computed the neighborhood graph of cells using the sc.pp.neighbors function with 10 nearest neighbors and 20 principal components. Clustering was performed using the Leiden algorithm (sc.tl.leiden) to assign cells to discrete clusters. To better understand the global connectivity structure of the data, we constructed a partition-based graph abstraction (PAGA) using the sc.tl.paga function, followed by visualization of the PAGA graph using UMAP initialized with PAGA positions (sc.tl.umap). UMAP plots were generated to visualize the clustering structure, with cells colored by Leiden clusters.

We log-transformed the counts using the sc.pp.log1p function and performed principal component analysis (PCA) using the sc.tl.pca function, with the ARPACK solver, to reduce the dimensionality of the dataset. We extracted the first two principal components (PCs) and calculated the variance ratio for each PC to assess the variance explained by each component. To further explore differences between clusters, we computed a distance matrix of the bulk data and performed hierarchical clustering to visualize these differences using single-linkage clustering. We visualized the hierarchical relationships between clusters with the scipy.cluster.hierarchy.dendrogram function. We calculated descriptive statistics, including the mean, median, standard deviation, and range of distances, to quantify the variability between clusters. To identify marker genes for each cluster, we performed differential expression analysis using the Wilcoxon rank-sum test (sc.tl.rank_genes_groups), chosen to account for the non-normal distribution of bulk RNA sequencing data.

We compared our dataset to three reference atlases: the Fly Cell Atlas, specifically the 10X VSN All (Stringent) dataset without blood, which contains whole transcriptome data from 5-day-old *D. melanogaster* nuclei generated using the 10X Genomics platform (Li et al. 2022); an Embryo Atlas, which provides temporal resolution of *Drosophila* embryonic development (Peng et al. 2024); and a Myeloid Blood Cell Atlas, which offers detailed characterization of Drosophila blood cell lineages (Cho et al. 2020). For each atlas, we imported the data into Scanpy (Wolf et al. 2018) and filtered the datasets: cells expressing fewer than 200 genes were filtered out, and genes expressed in fewer than three cells were removed using the sc.pp.filter_cells and sc.pp.filter_genes functions. The filtered data was normalized to a target sum of 1e4 counts per cell (sc.pp.normalize_total).

To ensure balanced representation of cell types for reference mapping, we implemented a subsampling strategy where each cell type was represented by an equal number of cells, determined by the size of the smallest cell type group. This was accomplished using custom subsampling functions that randomly selected cells from each category while maintaining the overall cell type distribution.

To integrate our bulk RNA-seq data with each single-cell reference atlas, we performed batch correction to account for technical differences between the datasets. We first subset both datasets to include only shared genes, which ensured comparable feature spaces. The datasets were then concatenated with appropriate batch labeling (bulk_adata.obs['dataset'] = 'Bulk', ref_adata.obs['dataset'] = 'Single-cell'). We applied the BBKNN algorithm (Polański et al. 2020a) for batch effect correction with the 'dataset' variable as the batch key.

For classification of bulk samples, we implemented a k-nearest neighbors (kNN) approach that assigned cell type labels based on proximity in the reduced dimensional space. The optimal k value was determined algorithmically as the minimum of: 1) the square root of the reference cell count and 2) 10% of the smallest cell type class, with a minimum threshold of 5 neighbors. This approach balanced statistical power with the risk of overfitting. For each bulk sample, we identified its k nearest reference cells and assigned a cell type based on a distance-weighted voting system where closer neighbors had greater influence on the classification outcome.

To assess classification confidence, we calculated a weighted score based on the proportion of neighbors belonging to the assigned cell type, with distances used as weights. We also implemented a permutation test (1000 iterations) to calculate p-values for each assignment, where labels were randomly shuffled to determine the likelihood of obtaining the observed classification by chance.

For visualization, we computed principal components using the sc.tl.pca function with the ARPACK solver, followed by neighborhood graph construction (sc.pp.neighbors) using 15 neighbors and 30 principal components. UMAP dimensionality reduction was applied to the integrated data (sc.tl.umap), allowing visualization of bulk samples within the context of reference cell types. We generated enhanced visualizations highlighting both experimental conditions (JW18 uninfected, JW18 *w*Mel, S2 uninfected, S2 *w*Mel) and predicted cell type assignments across each of the three reference atlases.

e. *Differential expression*

Following short-read processing and pseudoalignment, we filtered out transcripts with less than 10 reads across all replicates of each condition. This resulted in the detection of 10,839 expressed transcripts, out of 35,344 transcripts in the *D. melanogaster* genome and 1,122 expressed transcripts, out of 1,286 transcripts in the *w*Mel genome. We performed Wald tests to detect differential expression while accounting for multiple testing by calculating FDR/Benjamini-Hochberg p-value corrections in DESeq2 (Love et al. 2014). For these tests, we modeled the impact of the experimental conditions on the Drosophila transcriptomes as a function of cell line, infection, and the interaction between genotype and infection (~cell line + infection + cell line*infection). *Wolbachia* transcriptomes were tested for an impact of cell type on expression (~cell line).

f. *Gene set enrichment analysis (GSEA)*

For each set of differentially expressed genes, we imported the DEseq results into R, ranked the expressed genes by stat value, from high to low order, and performed GSEA with the clusterProfiler package (Yu et al. 2012). The stat value was selected because it incorporates information from both the adjusted-p value and the log2-fold change value. We performed KEGG enrichment analysis on the DE hits with gseKEGG (organism = "dme", minGSSize = 10, maxGSSize = 500, pvalueCutoff = 0.05). After mapping the FLYBASECGs ids to the

ENTREZIDs with the "org.Dm.eg.db" database, we performed GO enrichment analysis with gseGO (minGSSize = 10, maxGSSize = 500, pvalueCutoff = 0.05).

g. *Weighted gene co-expression network analysis (WGCNA)*

For each transcriptome, we filtered counts to only include genes with greater than 70 reads across all replicates of each condition, resulting in 8,322 expressed *D. melanogaster* transcripts and 763 *w*Mel transcripts, which were then normalized for composition and transformed with the regularized log function, rlog. We used the pickSoftThreshold R function (Horvath and Dong 2008) to find the best fit scale free topology value for network construction, which was 18 for *D. melanogaster* and eight for *w*Mel. Using the blockwiseModules function in R (Zhang and Horvath 2005), we clustered genes and samples hierarchically using signed networks and signed TOMtypes, Pearson correlation, and automated block-wise clustering to group associated genes based on their expression patterns into eigengene modules. Clustering resulted in 13 *D. melanogaster* modules and five *w*Mel modules. We tested the eigengene modules for association with the terms of the differential expression model ~ cell type + infection + cell type * infection by linear regression with limma::lmFit in R (Newville et al. 2014). We applied empirical Bayes, limma::eBayes, to smooth standard errors.

h. *Enrichment Analysis*

We performed gene ontology (GO) enrichment analysis on the DE genes in each category to identify pathways that are overrepresented compared to the background transcriptome (i.e., the full sets of expressed transcripts). Lists of genes comprising each eigengene module were compared with the background expressed reference transcriptome sets of *D. melanogaster* and *w*Mel gene IDs with clusterProfiler (Wu et al. 2021; Xu et al. 2024; Yu et al. 2012) and ShinyGO (Ge et al. 2020). *D. melanogaster* gene sets were also analyzed with the *Drosophila*-specific tool, PANGEA (Hu et al. 2023).

6. <u>*Drosophila* cell atlas *Wolbachia* titer estimation</u>

We investigated the *Wolbachia* infection status of three Drosophila cell atlas datasets (Fly Cell Atlas, Embryo Cell Atlas, and Myeloid Cell Atlas), by comparing against known infected and uninfected scRNAseq controls obtained from testis data in SRA: PRJNA788731 (Dou et al. 2023). Reads were first processed with Trimmomatic (v0.39) to remove adapter sequences and low-quality bases prior to alignment. We aligned transcriptomic data from each atlas to a combined reference genome containing both *Drosophila melanogaster* and *Wolbachia* *w*Mel sequences using STAR (v2.7.9a). For the cell atlas datasets, we generated subsampled data using seqtk (v1.4; https://github.com/lh3/seqtk), randomly sampling 1,000,000 reads from each SRA accession to ensure consistent depth across samples. We quantified gene expression using featureCounts from the Subread package. We extracted reads mapping to ribosomal RNA genes using established reference lists for both host and symbiont.

To estimate infection status of the scRNAseq atlases, we normalized the ratio of *w*Mel ribosomal RNA (rRNA) read counts to *D. melanogaster* rRNA counts for each sample. This approach takes advantage of mis-priming events of the poly-T primers used in scRNAseq to regions of ribosomal RNA enriched in adenine. We normalized read counts by transcript length to account for gene size differences. We employed Fisher's exact test on contingency tables of normalized read counts to compare each test sample against known infected and uninfected controls. A sample was classified as "uninfected" if it had an rRNA ratio profile that was not significantly different from the uninfected control, but significantly different from the infected control ($p < 0.05$). Conversely, a sample was classified as "infected" if the pattern was reversed. Samples with consistent signals from both controls were labeled as "inconclusive," with the final determination based on the relative similarity of their ratios to each control.

We implemented a bootstrap analysis to estimate 95% confidence intervals for *w*Mel/*D. melanogaster* rRNA ratios. The distribution of rRNA read counts normalized by transcript length was visualized using scatter plots with logarithmic scaling with custom Python scripts. We also generated bar plots displaying sample ratios with their associated 95% confidence intervals to facilitate direct comparison between samples.

7. Micro-C Chromatin Conformation Capture

To understand shifts in chromatin structure within infected cell lines we conducted a chromatin capture assay using the Dovetail Genomics Micro-C kit (Dovetail #: 21006). Micro-C is a nuclease-based proximity ligation method developed in 2015, derived from the original 3C method developed in 2002 (Dekker et al. 2002; Lee et al. 2022). The unique MNase used in Micro-C allows for high-resolution analysis of chromatin contacts while maintaining the ability to identify long-range interactions (Goel and Hansen 2021; Hsieh et al. 2020). These interactions include interactions within 1kb, such as enhancer-promoter loops, and long-range interactions, such as topologically associated domains (TADs) which can span lengths greater than 1Mb (Hsieh et al. 2020, 2022). This broad range makes the Micro-C method ideal for characterizing *Wolbachia* infections as the effects on chromatin structure have not previously been identified. We performed a Micro-C chromatin capture assay using the materials and methods from Dovetail's Micro-C Kit (Dovetail #: 21006) followed by an Illumina library preparation on two biological replicates per condition. Unless specified all materials were obtained from Dovetail Genomics.

a. *Cell culture and Crosslinking*

We collected immortalized *D. melanogaster* cell culture cells stably infected with *Wolbachia* from confluent cultures grown at 23°C. For each sample, 1 mL (at ~2e6 cells/mL) of cells were pelleted by centrifugation at 3,000xg for 5 minutes at 4°C. The supernatant was then removed and we resuspended cells in 1X PBS. We then repeated the centrifugation step and removed the PBS. Following supernatant removal, we promptly transferred the cell pellets to -80°C for overnight storage.

We performed a dual crosslinking protocol with formaldehyde and DSG (disuccinimidyl glutarate) to fix protein-DNA and protein-protein interactions. Cell pellets were thawed on at room temperature and resuspended in 1mL 1X PBS, 0.3 mM DSG each followed by a 10 minute rotation at room temperature on a hula mixer. We then added 27 µl of 37% formaldehyde to each sample followed by another 10 minute rotation. The samples were then pelleted by centrifugation at 3,000 x g for 5 minutes and washed twice with 1X wash buffer followed by centrifugation.

b. *Digestion and Lysis*

The pelleted material was resuspended in 50 μL of freshly prepared 1X Nuclease Digest Buffer and 0.5 μL of MNase Enzyme mix to fragment the crosslinked chromatin. Following crosslinking, we proceeded with the Dovetail Micro-C chromatin capture assay kit (Dovetail #: 21006) protocol and materials.

We performed a shallow sequencing run to determine the quality of the libraries (Supplemental Table S1) before full-depth sequencing. Paired-end Illumina reads were generated following the NEBNext Illumina library protocol. Libraries were sequenced on a NovaSeq S4 2x150bp lane by Fulgent Genetics. The demultiplexed sequencing reads underwent initial processing with Trimmomatic (v0.39) to process paired-end Illumina sequencing data with Phred+64 quality encoding, trim Illumina adapters, perform quality trimming by removing leading and trailing bases with Phred scores below 3, and discard reads shorter than 36 bases (Bolger et al. 2014).

c. *Mapping and pairing Micro-C contacts*

We processed these trimmed reads following recommendations from Dovetail Genomics (https://micro-c.readthedocs.io/). Briefly, we aligned the reads to the *D. melanogaster* genome *D. melanogaster* reference genome (GCF_000001215.4_Release_6_plus_ISO1_MT) with the Burrows-Wheeler Alignment tool (BWA, v0.7.17) (Li 2013a) using the BWA-MEM algorithm with a minimum quality score of 0 without pairing. The alignment results were parsed using pairtools, aiming to identify ligation junctions with a minimum quality score of 40 and a walks policy of 5 unique to report the 5'-most unique alignment on each side. Pairtools was also used to eliminate optical duplicates and produce read pairs for subsequent analysis (Abdennur et al. 2024). Following this, pairix was used to index the paired reads, and cooler (v 0.9.3) (Abdennur and Mirny 2019) was employed to construct chromatin contact maps with a bin size of 1kb. Contact maps were balanced at multiple resolutions with the cooler zoomify –balance function. We plotted the chromatin contact map using the Python tool CoolBox (Xu et al. 2021).

*d. Differential chromatin interaction analysis*

We extracted chromatin contacts from multi-resolution .mcool files at four resolutions (1 kb, 8 kb, 32 kb, and 128 kb) using cooler v0.9.3 (Open2C et al. 2024). For each biological replicate, we extracted both cis (intrachromosomal) and trans (interchromosomal) interactions across all *D. melanogaster* chromosomes (2L, 2R, 3L, 3R, 4, X, Y) using cooler dump -t pixels --join. Cis interactions were extracted for each chromosome individually, while trans interactions were extracted for all pairwise chromosome combinations. Interactions with balanced contact values equal to zero or NaN were filtered out, and the resulting contact matrices were consolidated into a single file containing genomic coordinates (chromosome, start, end positions for both anchors), read counts, replicate information, condition, interaction type (cis/trans), and resolution.

We converted these contact data to InteractionSet objects and combined biological replicates for differential analysis using diffHic v1.38 (Lun and Smyth 2015). Interactions were filtered to retain only those present in at least two samples with a minimum count of five reads and a minimum total count across all four samples of 20 reads. Count matrices were normalized using TMM normalization in edgeR v4.4.0 (Robinson et al. 2010), and dispersions were estimated using estimateDisp with robust=TRUE. For datasets exceeding 100,000 interactions, dispersion was first estimated on a random subset of 50,000 interactions and then applied to the full dataset to improve computational efficiency. Differential interactions were identified using glmQLFit and glmQLFTest for smaller datasets or glmFit and glmLRT for datasets exceeding 500,000 interactions. Statistical significance was assessed at FDR < 0.05 using the Benjamini-Hochberg method (Benjamini and Hochberg 1995), with genomic distances calculated for cis interactions. A likelihood ratio test comparing the full model (~infection) to a null intercept-only model was performed to assess overall significance of infection status on chromatin architecture.

*e. Identification and filtering of contacts within structural variants*

Cell lines can accumulate structural variants (SVs) relative to reference genomes, potentially confounding chromatin conformation analyses by creating artifactual differences in contact mapping (Cubeñas-Potts et al. 2017). To address this concern, we sequenced JW18 genomic DNA using Oxford Nanopore long-read technology and identified structural variants using SVIM (Heller and Vingron 2019) and removed any differential chromatin interactions overlapping SV breakpoints from subsequent analyses.

To generate long read data, wMel-infected JW18 cells, 1.2 mL (at ~ 2e6 cells/mL) of cells were pelleted by centrifugation at 16,000xg for 10 min at 4°C. We removed the supernatant and, extracted DNA using the Wizard HMW DNA Extraction kit (Promega #A2920). We prepared libraries with the Native Barcoding Kit V14 for Nanopore MinION R10 (Oxford Nanopore Technologies Cat #SQK-NBD114-24) and sequenced the libraries with a Nanopore MinION Mk1B with a MinION R10 Version flow cell (FLO-MIN-114, Lot:11,003,064). We used Oxford Nanopore's MinKNOW v23.07.8 software to live basecall with Guppy v7.0.8 (Fast model, read splitting ON) with the minimum read length set to 200 bp. Sequencing was stopped after 36 hours resulting in 3.65 M reads with an estimated N50 of 1.11 kb and 2.6 Gb called with a minQ of 8.

We processed the raw nanopore reads to select for only host-derived sequences. Reads were aligned to the *D. melanogaster* reference genome (dmell-all-r6.46) with bwa-mem v0.7.17 (Li 2013b). We used samtools v1.6 (Li et al. 2009) to sort and index the alignment and output only reads which aligned to the host genome (samtools view -b -F 4). We exported these reads as fastq files with bedtools v2.31.1 bamtofastq (Quinlan and Hall 2010). Structural variants were identified using SVIM v2.0.0 (Heller and Vingron 2019) in nanopore mode (svim reads

--nanopore) against the *D. melanogaster* reference genome. We filtered the resulting variants for high-quality calls (QUAL ≥ 10) using bcftools v1.22 (Danecek et al. 2021). To ensure that differential chromatin interactions identified by diffHic were not confounded by structural variation, we removed diffHic calls that overlapped with SVIM-identified structural variant breakpoints.

SVIM identified 1485 deletions, 1014 insertions, 26 breakends, 59 tandem duplications, and one interspersed duplication. Chromatin interactions that either anchored to or overlapped with these SV regions (with 200 bp padding around each SV breakpoint) were removed from downstream analyses, along with *cis* interactions shorter than 4 kb to focus on long-range chromatin contacts. This approach ensures that observed differences in chromatin contacts reflect infection-mediated changes unrelated to underlying genomic structural variation between our cell line and the *D. melanogaster* reference genome (Ghavi-Helm et al. 2019; dos Santos et al. 2015). Filtering reduced interactions from 42.0M and 41.7M (uninfected) to 29.2M and 11.0M, and from 157.3M and 156.9M (*w*Mel-infected) to 107.1M and 102.8M, retaining 26-69% of interactions across replicates.

*f. A/B Compartment analysis*

To assess changes in higher-order chromatin organization, we performed A/B compartment analysis using cooltools v0.7.4 (Open2C et al. 2024) across 1,851 genomic bins. Compartment analysis was performed on balanced contact matrices we extracted from multi-resolution .mcool files at 50 kb resolution. For each condition, we calculated the first three eigenvectors (E1, E2, E3) of the observed/expected contact matrix using cooltools.eigs_cis with default parameters (ignore_diags=2, clip_percentile=99.9). The sign of the first eigenvector (E1) was oriented such that positive values correspond to gene-dense, transcriptionally active A compartments and negative values correspond to gene-poor, transcriptionally inactive B compartments. Compartment assignments (A or B) were determined by the sign of E1 for each 50 kb genomic bin.

To identify regions with significant compartment changes between uninfected and infected conditions, we merged compartment data from both conditions based on genomic coordinates (chromosome, start, end positions). For each genomic bin, we calculated the E1 difference (E1_infected - E1_uninfected) and identified compartment switches (A→B or B→A) where the compartment label changed between conditions. To assess statistical significance, we calculated z-scores for the absolute E1 difference at each bin: $z = (|E1\_diff| - \mu) / \sigma$, where μ and σ are the genome-wide mean and standard deviation of |E1_diff|. Two-tailed p-values were calculated from z-scores using the standard normal cumulative distribution function and corrected for multiple testing using the Benjamini-Hochberg method (Benjamini and Hochberg 1995). Bins with FDR < 0.01 were considered to have significant compartment changes. Global significance of compartment changes was assessed using a paired t-test comparing E1 values between conditions across all genomic bins.

To assess whether the overall rate of compartment switching between conditions exceeded random expectation, we evaluated the observed switch rate against a label-permutation null model. Compartment labels (A or B) were randomly permuted between conditions 1,000 times to generate a null distribution of switch rates, from which empirical two-sided p-values were derived (as in (Ghavi-Helm et al. 2019)). For genotype-specific comparisons, we used Fisher's exact test to determine whether compartment switch rates differed significantly between bins with and without significant E1 changes. Chromosome-specific enrichment of compartment switches was evaluated by comparing the distribution of A→B versus B→A switches across major chromosome arms (2L, 2R, 3L, 3R, X).

g. *Topologically Associated Domains (TADs), Hotspots, and Loops*

We quantified insulation strength to map topologically associated domains (TADs) and chromatin loops using cooltools.insulation (Open2C et al. 2024) on contact matrices at 50 kb resolution. First, we calculated insulation scores representing the normalized number of contacts between a window upstream and downstream for a biological condition. To do this, we calculated TAD separation scores as the average Z-score of all Micro-C contacts between adjacent 150 kb diamond-sliding windows upstream and downstream. Then, we then averaged these scores into an aggregate insulation score profile for uninfected and infected JW18 conditions (as in (Ghavi-Helm et al. 2019)). Local minima in the insulation profile were identified to call standard TAD boundaries for each condition. Second, boundary strength was calculated from the peak prominence to the minima of each insulation profile. Significant TADs were called after thresholding on boundary strength with the Li method (default threshold="Li"). We called loops with cooltools.dots, using a max_loci_separation of 2000000, fdr=0.01, and clustering_radius of 20000.

To map regions undergoing significant local structural reorganization, we identified differential-interaction hotspots. Significant differential interactions (FDR < 0.05, |logFC| > 1) derived from diffHic were aggregated into 50 kb non-overlapping genomic bins. For each bin, we calculated a hotspot strength score by summing the absolute log2 fold-changes of all differential interactions anchoring within that window. Hotspots were then explicitly defined by peak calling on these binned distributions using scipy.signal.find_peaks (Virtanen et al. 2020). We compared hotspot strength between uninfected and infected conditions to determine if the observed changes in hotspot strength between conditions were statistically significant. We generated a null distribution by randomly permuting the condition labels of the hotspot strengths 1,000 times, allowing us to calculate an empirical two-tailed p-value for the structural differences.

h. *Enrichment analyses*

   i. *Insulator Protein Enrichment Analysis*

   To assess whether differential chromatin interactions were enriched at insulator binding sites, we obtained *D. melanogaster* insulator protein binding locations from modENCODE ChIP-seq data (Landt et al. 2012) and calculated contact-insulator overlap. We employed Schwartz et al.'s classification of Class I insulators

(CTCF-dependent) and Class II insulators (CTCF-independent, including Su(Hw), BEAF-32, CP190, and Mod(mdg4)) (Schwartz et al. 2012). Insulator binding sites were extended by ±10 kb to account for the influence range of insulator proteins on chromatin organization (Vorobyeva et al. 2024). We calculated observed insulator overlap by intersecting differential chromatin interaction anchors with extended insulator regions using BedTools v2.31.1 (Quinlan and Hall 2010). For each interaction, we determined whether either anchor (any anchor overlap) or both anchors (both anchors overlap) intersected with insulator sites, and we counted the total number of insulators overlapping each anchor. We also calculated the minimum distance from each anchor to the nearest insulator binding site.

To assess statistical significance of insulator enrichment, we employed two complementary approaches. First, we calculated a genomic background model that estimates the expected probability of insulator overlap based on genome-wide insulator density. Total insulator coverage was calculated by merging overlapping extended insulator regions using BedTools merge, and chromosome-specific insulator densities were computed by dividing total insulator coverage by chromosome size. For each interaction, we calculated the expected probability that each anchor would overlap an insulator (P = insulator density), and the expected probability that any anchor would overlap an insulator was calculated as P(anchor1) + P(anchor2) - P(anchor1) × P(anchor2). Statistical significance was assessed using a binomial test comparing observed overlap counts to expected counts across all interactions. Second, we performed permutation testing with 1,000 iterations to assess enrichment relative to random genomic locations. For each permutation, interaction anchor coordinates were randomly shuffled across the genome while preserving chromosome identity using BedTools shuffle with the *D. melanogaster* genome file (dm6). Overlap rates were calculated for each permuted dataset, generating a null distribution of expected overlap rates. Empirical p-values were calculated as the proportion of permutations with overlap rates equal to or greater than the observed rate. Enrichment was calculated as the ratio of observed to expected (median null) overlap rates, and z-scores were computed from the null distribution mean and standard deviation.

We analyzed insulator enrichment separately for Class I and Class II insulators, and for up-regulated versus down-regulated interactions (logFC > 0 vs. logFC < 0). We tested for direction-specific enrichment using chi-square contingency tests. All analyses were performed in Python v3.9 using pandas v1.5.3, numpy v1.24.3, scipy v1.10.1, and pybedtools v0.9.0.

*ii. Enhancer Enrichment Analysis*

Enhancer regions were defined using housekeeping and developmental core promoter enhancer peak sets from (Zabidi et al. 2015) (GEO: GSE57876). Separate peak files were obtained for S2 and ovarian somatic (OSC) cell lines for each enhancer class (housekeeping core promoter, hkCP; developmental core promoter, dCP). For each class, peak summit coordinates (column 2 of each peaks file) were extracted as single-base intervals, pooled across both cell lines, and merged using bedtools merge (Quinlan and Hall 2010) following coordinate sorting, yielding a non-redundant set of housekeeping and developmental enhancer regions. Class labels were appended as a seventh column to produce a seven-column BED file, and "chr" prefixes were stripped from chromosome names to match the dm6 reference annotation used in chromatin contact analysis. The final combined enhancer reference contained 11,924 housekeeping and 11,136 developmental elements.

We intersected the enhancer reference set with the differential chromatin contacts between *w*Mel-infected JW18 cells and the uninfected DOX reference condition using pybedtools (Dale et al. 2011) to characterize enhancer-specific chromatin reorganization. Each interaction anchor was evaluated for coordinate overlap with the enhancer set; an overlap was defined as any non-disjoint interval intersection (Rao et al. 2014). Interactions in which both anchors overlapped annotated enhancers were designated enhancer-enhancer (E-E) contacts; interactions in which only one anchor overlapped an enhancer were designated enhancer-to-promoter (E-TSS) contacts (Javierre et al. 2016). E-E contacts were further subclassified by the class identity of the two participating enhancers: same-class contacts (both housekeeping or both developmental) were assigned accordingly (Rubin et al. 2017), while contacts bridging enhancers of different classes were designated cross-class.

To determine whether housekeeping and developmental enhancers responded differentially to *w*Mel infection, logFC distributions were compared pairwise between all contact class groups within each interaction type (E-E, E-TSS) using Welch's two-sample t-test and the Mann-Whitney U test (two-sided) as implemented in SciPy (Virtanen et al. 2020). Comparisons were restricted to groups with at least ten observations. Effect sizes were estimated using Cohen's d, interpreted as negligible ($|d| < 0.2$), small (0.2 to 0.5), medium (0.5 to 0.8), or large ($|d| > 0.8$). Bootstrap 95% confidence intervals for mean logFC differences were estimated from 1,000 resampling iterations with replacement. P-values from both tests were corrected for multiple comparisons using the Benjamini-Hochberg procedure, with statistical significance defined at FDR < 0.05. A secondary threshold of |mean logFC difference| > 0.5 was applied to distinguish effects of potential biological relevance from statistically significant but negligible differences.

*iii.* *Histone Modification Enrichment Analysis at X Chromosome Contacts*

We obtained H3K36me3 ChIP-seq peak coordinates from modENCODE for *D. melanogaster* Kc167 cells (modENCODE, GSE20784) and H4K16ac ChIP-seq peak coordinates from *D. melanogaster* embryos (Samata et al. 2020). H3K36me3 peak coordinates were converted from dm3 to dm6 genome assembly by the FlyBase

Coordinate Converter tool (https://flybase.org/convert/coordinates). We filtered peak files in BED format (chromosome, start, end, score) to retain only X chromosome peaks. Differential chromatin interactions were filtered to retain only those involving the X chromosome and meeting significance thresholds (FDR < 0.05, |logFC| > 1). For each histone modification, we extended interaction anchors by ±5 kb or ±50 bp respectively for H3K36me3 or H4K16ac sites to account for the spread of histone marks around contact sites (Gelbart and Kuroda 2009; Larschan et al. 2007). Overlap between extended interaction anchors and histone modification peaks was calculated using BedTools v2.31.1 (Quinlan and Hall 2010). For each interaction, we determined whether either anchor (any anchor overlap) or both anchors (both anchors overlap) intersected with histone modification peaks, and we counted the total number of peaks overlapping each anchor.

We assessed statistical significance of histone modification enrichment using a permutation test (n = 100 permutations). For each permutation, we randomly shuffled the genomic positions of interaction anchors across the X chromosome while maintaining their original sizes, then recalculated overlap with histone modification peaks. This approach controls the genomic distribution of histone marks, anchor sizes, and chromosome-specific features. Empirical p-values were calculated as twice the minimum of the proportion of permuted overlap rates greater than or equal to (or less than or equal to) the observed rate, providing a two-tailed test. Enrichment z-scores were calculated as (observed - null mean) / null standard deviation.

To test for genotype-specific effects, we analyzed histone modification enrichment separately for up-regulated interactions (logFC > 0, representing stronger contacts in uninfected JW18 cells) and down-regulated interactions (logFC < 0, representing stronger contacts in *w*Mel-infected JW18 cells). Differences in overlap rates between genotypes were tested using Fisher's exact test. For interactions classified as cis (intrachromosomal) or trans (interchromosomal), we compared histone modification enrichment using Fisher's exact test. Distributions of peak counts between genotypes were compared using the Mann-Whitney U test. All analyses were performed in Python v3.9 using pandas v1.5.3, numpy v1.24.3, scipy v1.10.1, and pybedtools v0.9.0. Visualizations were generated using matplotlib v3.7.1 and seaborn v0.12.2, with boxplots displaying the median, quartiles, and individual data points with jitter to reveal underlying distributions.

iv. *High-Affinity Site (HAS) Enrichment Analysis*

To assess whether differential chromatin interactions on the X chromosome were enriched at dosage compensation complex entry sites, we obtained high-affinity site (HAS), also called chromatin entry site (CES), coordinates from modENCODE ChIP-seq data for MSL (Male-Specific Lethal) complex binding (Alekseyenko et al. 2008; Straub et al. 2008). HAS/CES represent specific DNA sequence motifs where the MSL dosage compensation complex initially binds before spreading along the X chromosome to upregulate gene expression (Villa et al. 2016). We filtered differential chromatin interactions to retain only those involving the X chromosome. HAS/CES binding sites were extended by ±50 kb to account for the spreading range of the MSL complex from entry sites (Alekseyenko et al. 2008). We calculated the overlap rate between differential interaction anchors and extended HAS/CES regions using BedTools v2.31.1 (Quinlan and Hall 2010). For each interaction, we determined whether either anchor (any anchor overlap) or both anchors (both anchors overlap) intersected with HAS/CES sites, counted the total number of HAS/CES overlapping each anchor, and calculated the minimum distance from each anchor to the nearest HAS/CES site.

We assessed statistical significance of HAS/CES-contact enrichment using a binomial test comparing observed overlap rates to a null expectation of 7.5%, derived from the genomic coverage of HAS/CES sites with 50 kb extensions on the ~23 Mb X chromosome. Enrichment was calculated as the ratio of observed to expected overlap rates. To test for genotype-specific effects, we analyzed HAS/CES enrichment separately for up-regulated interactions (logFC > 0, representing stronger contacts in uninfected JW18 cells) and down-regulated interactions (logFC < 0, representing stronger contacts in *w*Mel-infected JW18 cells). Differences in overlap rates between genotypes were tested using chi-square contingency tests. Distance distributions to the nearest HAS/CES site were compared between genotypes using the Mann-Whitney U test. Spearman correlation was calculated between interaction effect size (logFC) and distance to the nearest HAS/CES site. All analyses were performed in Python v3.9 using pandas v1.5.3, numpy v1.24.3, scipy v1.10.1, and pybedtools v0.9.0.

8. Cell fixation and immunocytochemistry

The day before preparation, we seeded cells on no 1.5 glass coverslips at the bottom of 6-well plates (Corning 3516). Following removal of cell culture medium, we fixed the cells in 8% paraformaldehyde in 1xPBS for 12 minutes at room temperature. After washing three times in PBS-T, we blocked the cells in 1% bovine serum albumin in PBS-T for one hour at room temperature, and then incubated the cells in the primary antibodies diluted in PBS-T for an hour at room temperature (antibodies and dilutions listed below). Then, we washed the cells three times in PBS-T and incubated them in secondary antibodies diluted 1:500 in PBS-T for an hour at room temperature. We used mounting media containing DAPI (VECTASHIELD Antifade Mounting Medium With DAPI H-1200) to counterstain the cells. Primary monoclonal antibodies were used at the following dilutions in PBS-T: anti-phHis2Av at 1:500 (UNC93-5.2.1, Developmental Studies Hybridoma Bank (University of Iowa)) and anti-H4K16ac at 1:100 (600-401-J00, Rockland). Cells were imaged on a Leica Stellaris Confocal microscope with a 63x objective. Optical sections were taken at the Nyquist value for the objective. Images were processed in ImageJ.

**RESOURCES TABLE**

| REAGENT or RESOURCE | SOURCE | IDENTIFIER |
|---|---|---|
| **Bacterial and Virus Strains** | | |
| *Wolbachia w*Mel | Growing in *Wolbachia*-infected JW18 cells | |
| **Critical Commercial Assays** | | |
| FastSelect - rRNA Fly | Qiagen | Cat# 333262 |
| RNeasy Mini Kit | Qiagen | Cat# 74104 |
| Micro-C Kit | Dovetail | Cat# 21006 |
| Qubit dsDNA HS Kit | Invitrogen | Cat# Q33231 |
| SPRI-select beads | Beckman | Cat# B23318 |
| DNA Clean and Concentrator-5 Kit | Zymo Research | Cat# D4004 |
| NEBNext Illumina library | New England Biolabs | Cat# E7645L |
| Antifade Mounting Medium With DAPI | VECTASHIELD | Cat# H-1200 |
| **Deposited Data** | | |
| RNA-seq reads | This paper | PRJNA1240446 |
| Micro-C reads | This paper | PRJNA1240446 |
| **Experimental Models: Cell Lines** | | |
| JW18 | (Serbus et al. 2012) | FlyBase ID: FBtc9000069 |
| S2 | (Schneider 1972) | FlyBase ID: FBtc0000150 |
| **Software and Algorithms** | | |
| bwa (version 0.7.17) | (Li 2013b) | https://github.com/lh3/bwa |
| samtools (version 1.18) | (Li et al. 2009) | http://www.htslib.org/ |
| Pairtools (version 1.0.2) | (Abdennur et al. 2024) | https://pairtools.readthedocs.io/ |
| CoolBox (version 0.3.8) | (Xu et al. 2021) | https://gangcaolab.github.io/CoolBox/quick_start_API.html |
| Cooler (version 0.9.3) | (Abdennur and Mirny 2019) | https://cooler.readthedocs.io/en/latest/ |
| Mustache (version 1.0.1) | (Roayaei Ardakany et al. 2020) | https://github.com/ay-lab/mustache |
| Scanpy (version 1.9.3) | (Wolf et al. 2018) | https://scanpy.readthedocs.io/en/stable/ |
| Anndata (version 0.10.7) | (Virshup et al. 2024) | https://anndata.readthedocs.io/en/stable/ |
| Pandas (version 2.2.2) | The Pandas Development Team | https://pandas.pydata.org/ |
| Numpy (version 1.26.4) | (Harris et al. 2020) | https://numpy.org/ |
| BBKNN (version 1.6.0) | (Polański et al. 2020b) | https://github.com/Teichlab/bbknn |
| Scipy (version 1.13.0) | (Virtanen et al. 2020) | https://scipy.org/ |
| Matplotlib (version 3.8.4) | (Hunter 2007) | https://matplotlib.org/ |
| DESeq2 (version 1.40.2) | (Love et al. 2014) | https://bioconductor.org/packages/release/bioc/html/DESeq2.html |

| | | |
|---|---|---|
| trimmomatic (version 0.39) | (Bolger et al. 2014) | https://github.com/timflutre/trimmomatic |
| Tximport (version 1.28.0) | (Soneson et al. 2016) | https://bioconductor.org/packages/release/bioc/html/tximport.html |
| kallisto (version 0.48.0) | (Bray et al. 2016) | https://github.com/pachterlab/kallisto |
| python (version 3.10.12) | Python | https://www.python.org/ |
| R (version 4.3.1) | R-Project | https://www.r-project.org/ |
| snakemake (version 7.32.4) | (Mölder et al. 2021) | https://snakemake.readthedocs.io/en/stable/ |
| PANGEA (version 1.1) | (Hu et al. 2023) | https://www.flyrnai.org/tools/pangea/web/home/7227 |
| WGCNA (version 1.73) | (Langfelder and Horvath 2008) | https://cran.r-project.org/web/packages/WGCNA/index.html |
| Dplyr (version 1.1.4) | (Wickham et al. 2019) | https://dplyr.tidyverse.org/ |
| Pheatmap (version 1.0.12) | (Kolde 2025) | https://github.com/raivokolde/pheatmap |
| Genefilter (version 1.88.0) | (Gentleman et al. 2025) | https://bioconductor.org/packages/release/bioc/html/genefilter.html |
| Rhdf5 (version 2.50.2) | (Fischer et al. 2025) | https://www.bioconductor.org/packages/release/bioc/html/rhdf5.html |
| Ggforce (version 0.5.0) | (Pedersen and RStudio 2025) | https://ggforce.data-imaginist.com/ |
| ComplexHeatmap (version 2.23.0) | (Gu et al. 2016) | https://github.com/jokergoo/ComplexHeatmap |
| Tidyr (version 1.3.1) | (Wickham et al. 2019) | https://tidyr.tidyverse.org/ |
| Readr (version 2.1.5) | (Wickham et al. 2019) | https://readr.tidyverse.org/ |
| flashClust (version 1.01-2) | (team and Langfelder 2012) | https://cran.r-project.org/web/packages/flashClust/index.html |
| AnnotationDbi (version 1.68.0) | (Pagès et al. 2025) | https://bioconductor.org/packages/release/bioc/html/AnnotationDbi.html |
| clusterProfiler (version 4.14.6) | (Wu et al. 2021; Yu et al. 2012) | https://guangchuangyu.github.io/software/clusterProfiler/ |
| org.Dm.eg.db (version 3.20.0) | (Carlson 2025) | https://bioconductor.org/packages/release/data/annotation/html/org.Dm.eg.db.html |
| Enrichplot (version 1.18.3) | (Guangchuang and Chun-Hui 2025) | https://bioconductor.org/packages/release/bioc/html/enrichplot.html |
| Ggplot2 (version 3.5.1) | (Wickham et al. 2019) | URL:https://ggplot2.tidyverse.org/ |
| Scripts written for processing transcriptomic data | This paper | URL:https://github.com/shelbirussell/Dual_bulk_RNAseq_analysis-Jacobs_et_al.git |

| | | |
|---|---|---|
| Scripts written for processing Micro-C data | This paper | URL:https://github.com/jodiejacobs/wolbachia_induced_differentiation.git |
| **Other** | | |
| *Wolbachia w*Mel genome | (Wu et al. 2004) | GCF_000008025.1 |
| *Drosophila melanogaster* genome | (Hoskins et al. 2015) | GCF_000001215.4; Release_6_plus_ISO1_MT |
| Myeloid blood cell atlas | (Cho et al. 2020) | http://big.hanyang.ac.kr/flyscrna |
| Embryo Atlas | (Peng et al. 2024) | https://github.com/CahanLab/fruitfly_organogenesis |
| Fly CellAtlas (10x VSN All; Stringent) | (Li et al. 2022) | https://flycellatlas.org/ |

**Resource availability**
Further information and requests for resources and reagents should be directed to and will be fulfilled by Shelbi Russell.

**Materials availability**
This study did not generate new unique reagents. All cell lines can be obtained from the corresponding authors of the original publications (Mirchandani et al. 2024b; Serbus et al. 2012).
