## Supplemental Figures for "*Wolbachia* infection induces host cell state changes *in vitro* and determines symbiotic fate in *Drosophila*"

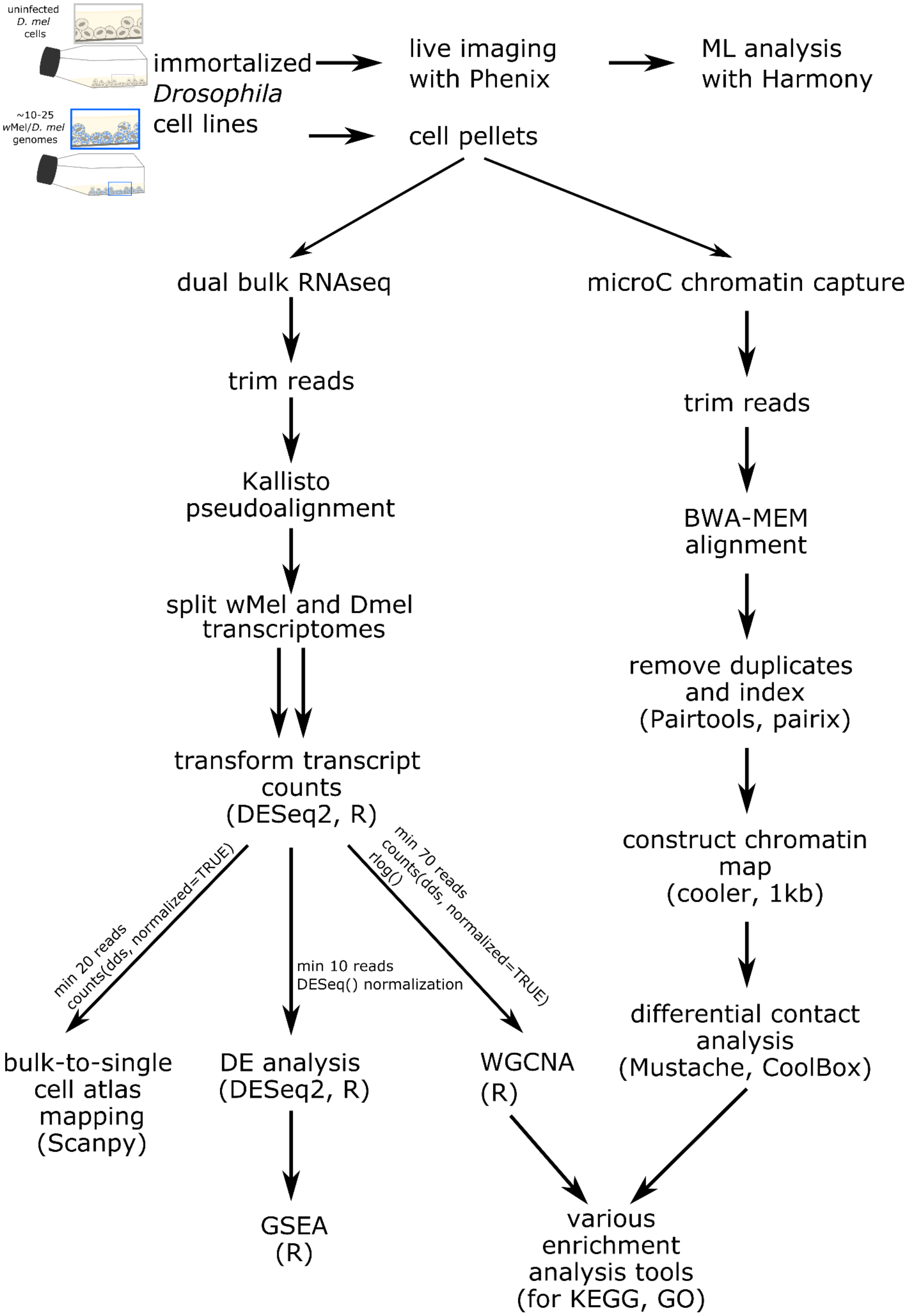


**Figure S1**. Methods overview

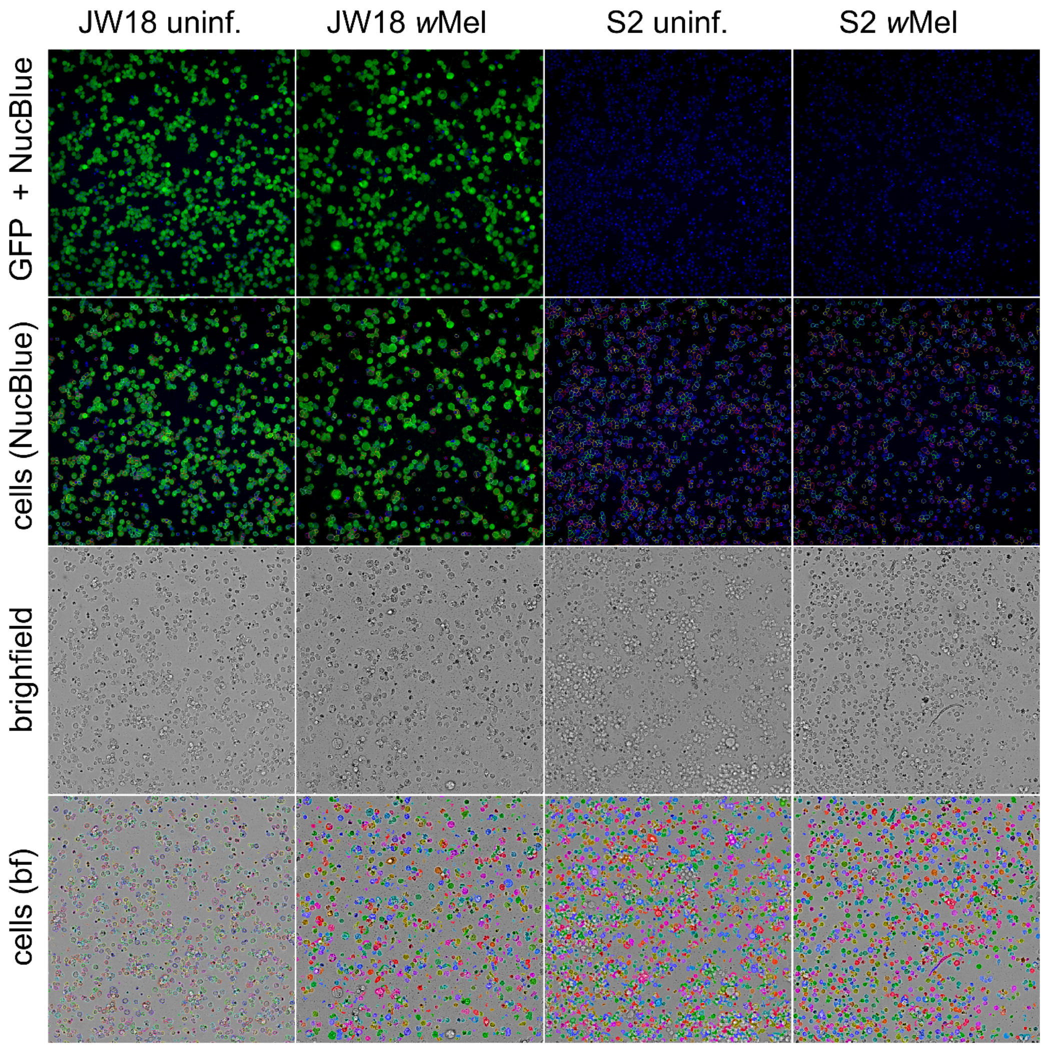


**Figure S2**. Representative Opera Phenix images from analysis in Fig. 2S-V.

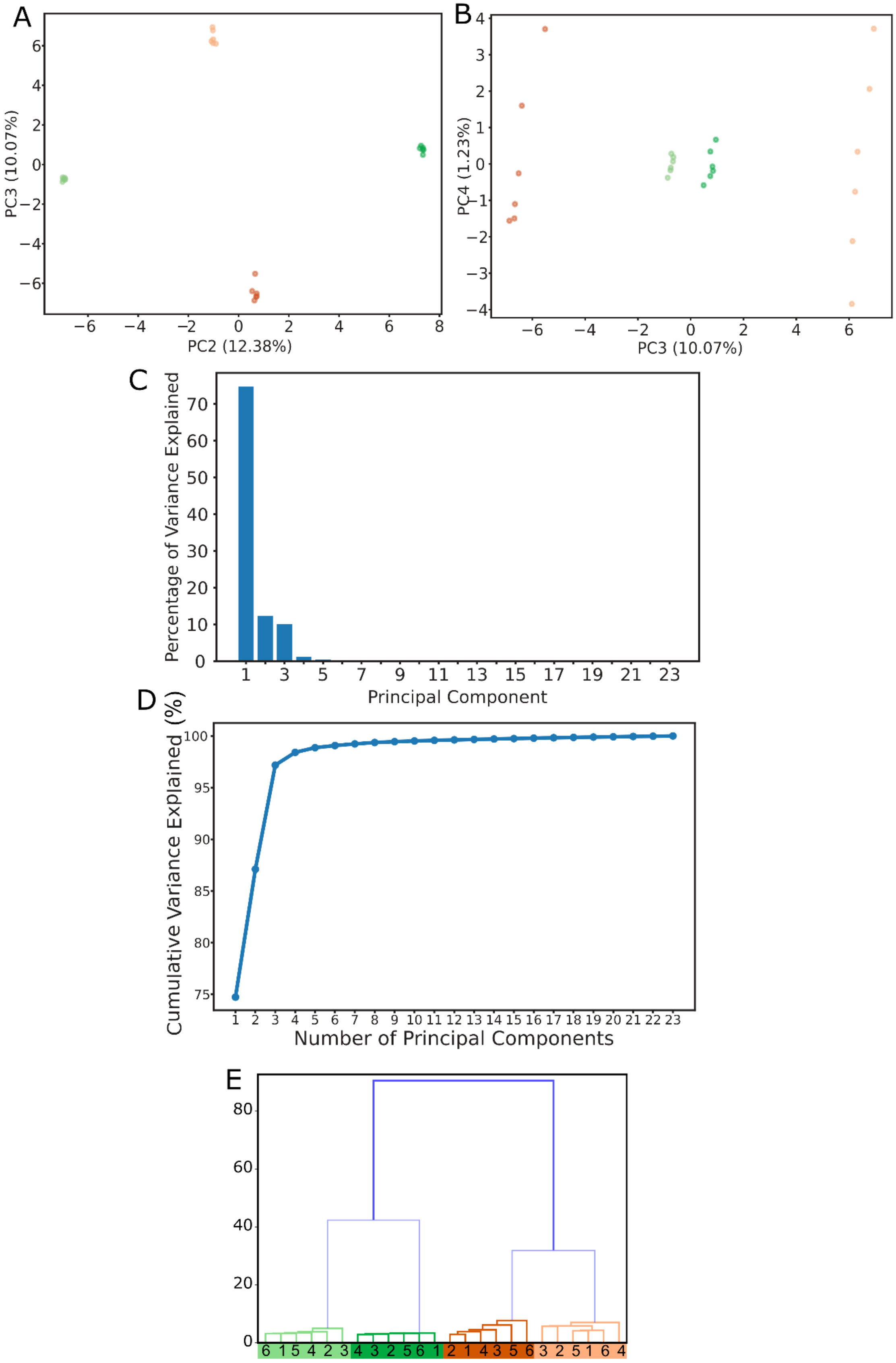


**Figure S3**. Bulk transcriptome PCA clustering results. A-B) PCA eigenvalue plots of PCs 2-4. C) Bar plot of explained variance by principal component. D) Cumulative explained variance as principal components are added. E) Hierarchical clustering dendrogram by dissimilarity, demonstrating primary separation by cell line, and secondary separation by *Wolbachia* infection status.

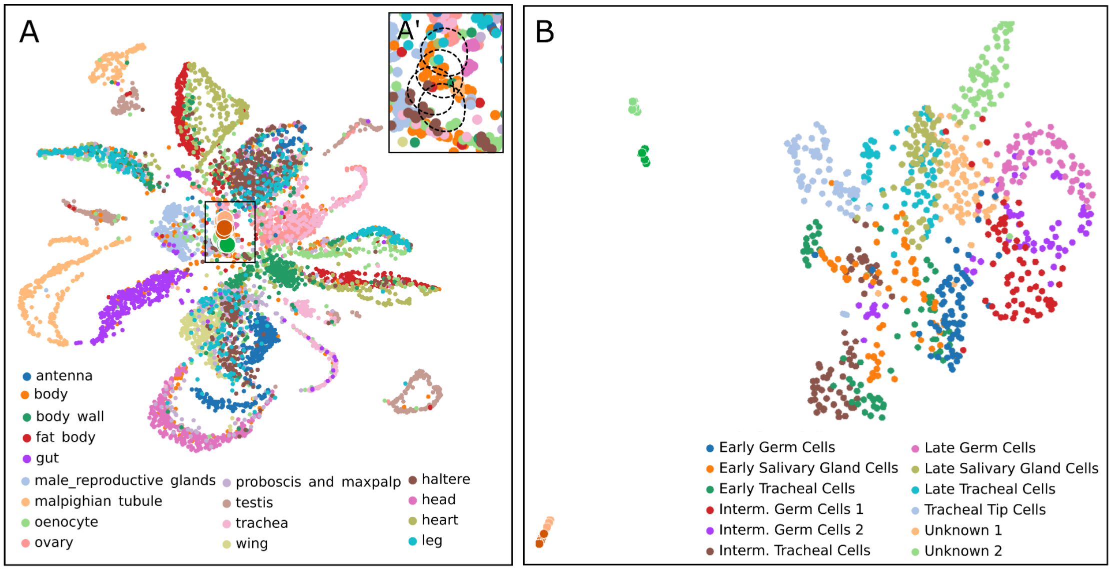


**Figure S4**. Mapping to the *Drosophila melanogaster* (A) adult and (B) embryo-larval cell atlas. A') Highlights where our bulk cell transcriptomes mapped to the adult atlas. UMAP projections of bulk RNA-seq data onto single-cell reference atlas for A) adult and B) embryonic *Drosophila*.

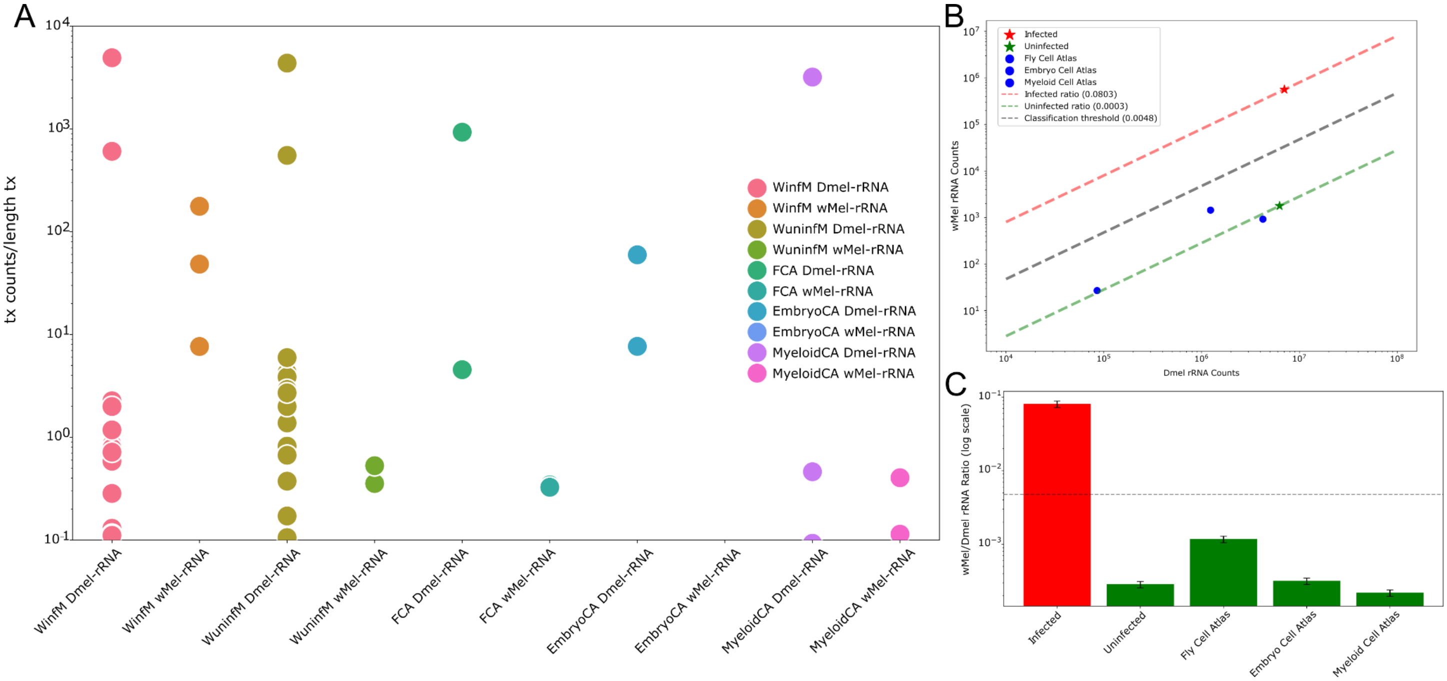


**Figure S5**. *Drosophila* atlas rRNA titers. A) An infected testis 3'-scRNAseq sample has higher numbers of reads mapping to the *w*Mel 16S rRNA than uninfected samples. Whereas, high numbers of reads map to *D. melanogaster* rRNAs in all samples. B) Comparing mapped *w*Mel rRNA counts to *D. melanogaster* rRNA counts indicated a detection threshold that can be used to call infected vs uninfected samples from poly-A-primed data. C) Relative *w*Mel:*D.melanogaster* rRNA coverage suggests that samples with *w*Mel:*D.mel* rRNA ratios less than 1e-2 are uninfected.

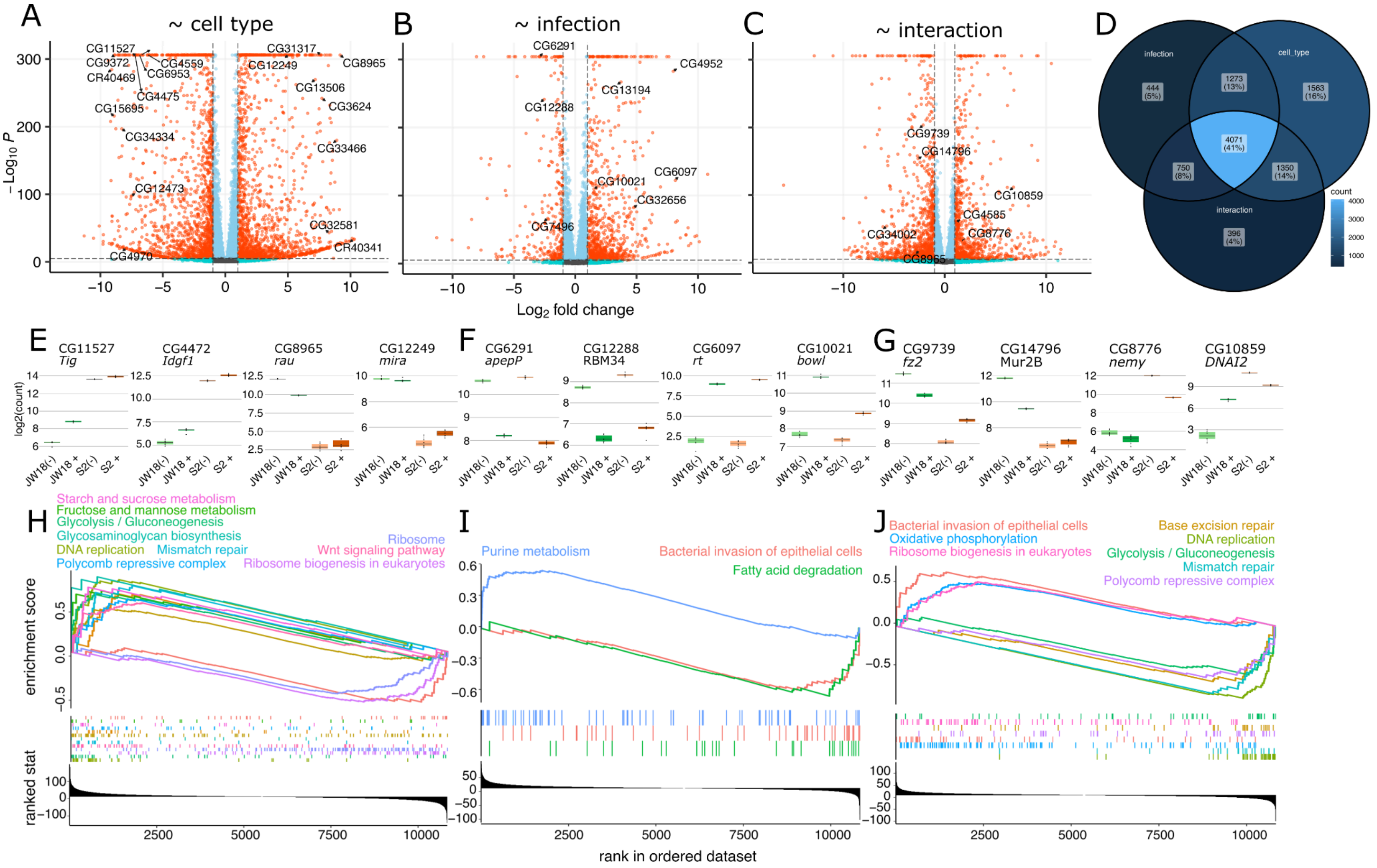


**Figure S6**. *D. melanogaster* bulk transcriptome DESeq2 differential expression analysis and Gene Set Enrichment Analysis (GSEA) results. A-C) DESeq2 differential expression results, plotted as volcano plots. Total genes expressed = 10,839; Colored points: non-significant=grey, Log2FC>1=teal, padj<1e-5=light blue, and Log2FC>1 and padj<1e-5=orange-red. D-F) Gene count plots D) Overlap among model terms for Wald Test differential expression hits. E-G) Gene count plots for some of the highly up and down-regulated genes labeled in the volcano plots. H-I) GSEA output ranked by the "stat" value output from DESeq2, which incorporates both the fold-change and adjusted p-value. H) Cell type GSEA: Gene ontology (GO) pathways significantly up- and downregulated in JW18 cells relative to S2 cells (p ≤ 0.05). I) Infection GSEA: GO pathways significantly up- and downregulated in uninfected cells relative to *w*Mel-infected cells (p ≤ 0.05). J) Interaction GSEA: GO pathways significantly up- and downregulated in JW18*w*Mel cells relative to S2 uninfected cells (p ≤ 0.05).

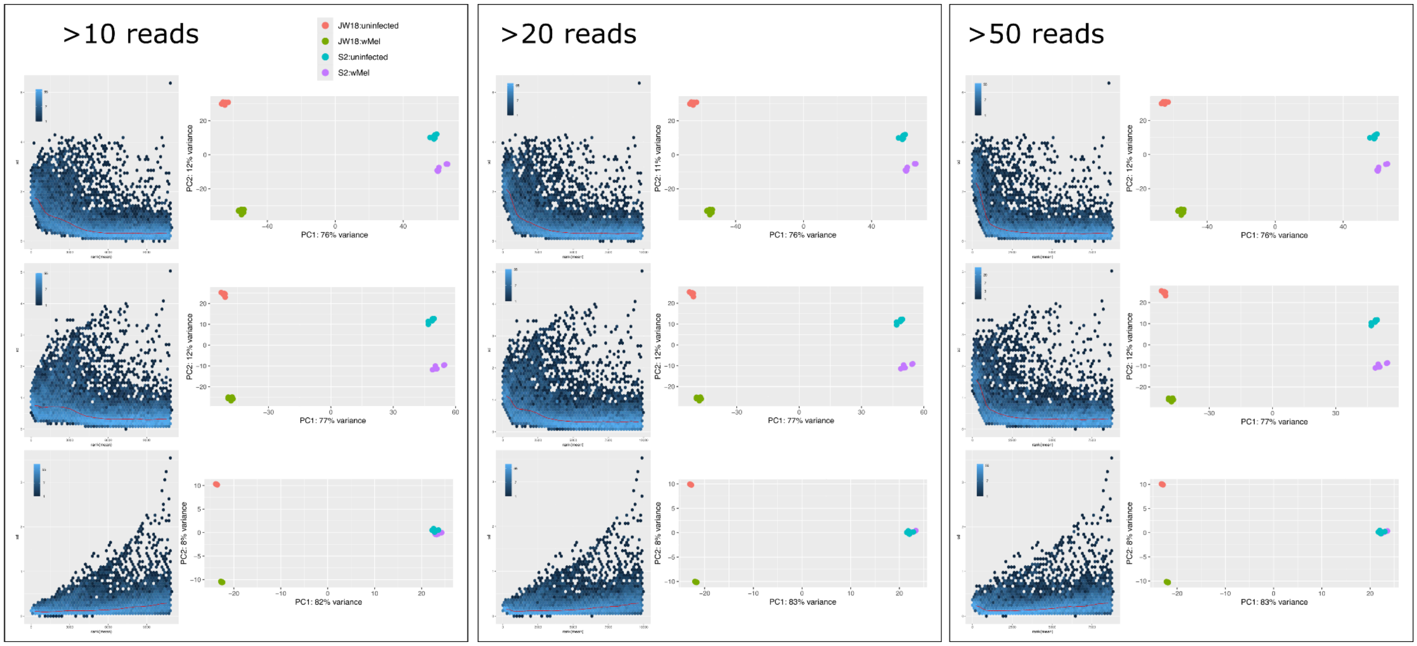


**Figure S7**. Transformation method selection for *Drosophila* WGCNA. We selected the regularized log (rld) transformation because the variance stabilizing (vsd) transformation removes infection-related variance in the S2 line.

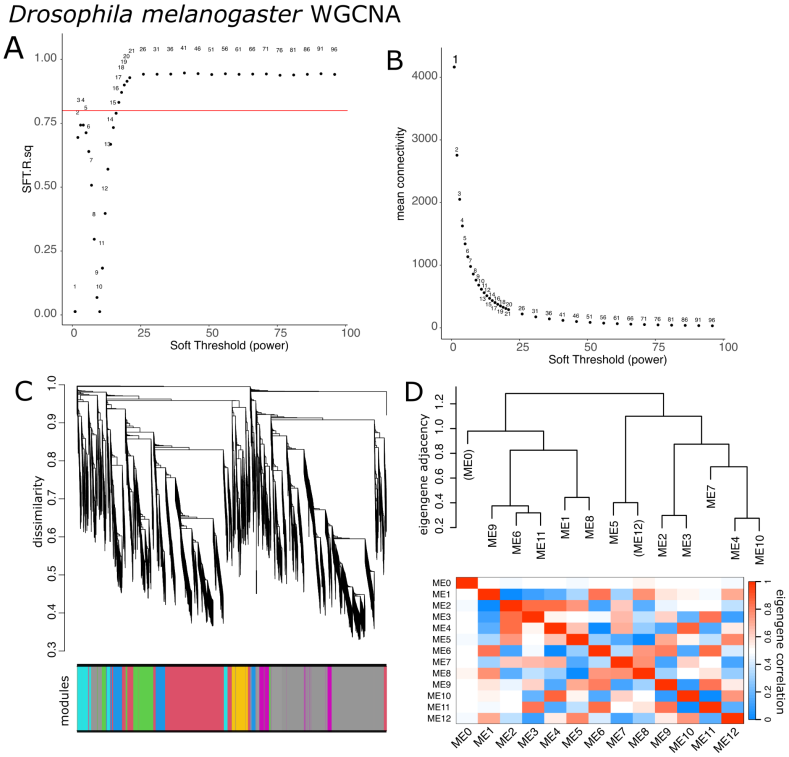


**Figure S8**. *Drosophila melanogaster* transcriptome WGCNA reveals expression modules associated with cell type, infection, and joint-cell type and infection states. A, B) Power thresholding analysis. C) Dendrogram and D) heatmap depicting relationships among the modules' eigengenes, the collapsed, combined, and normalized expression of the genes that make up each module.

A

| module | metadata. infectionwMel | metadata. celltypeS2 | AveExpr | F | P.Value | adj.P.Val | genes in module | DE padj>=0.01 genes in module | %DE padj>=0.01 genes in module |
|---|---|---|---|---|---|---|---|---|---|
| ME2 | -0.0311 | -0.4070 | -8.10E-18 | 3618.41 | 1.61E-28 | 2.09E-27 | 1952 | 1862 | 95.39% |
| ME9 | -0.3972 | 0.0744 | -5.55E-17 | 479.80 | 6.34E-19 | 4.12E-18 | 294 | 272 | 92.52% |
| ME1 | 0.0205 | 0.4020 | -2.66E-17 | 356.07 | 1.57E-17 | 6.79E-17 | 2687 | 2583 | 96.13% |
| ME4 | 0.2461 | -0.3035 | 1.62E-17 | 117.26 | 1.73E-12 | 5.61E-12 | 556 | 524 | 94.24% |
| ME6 | -0.2105 | 0.3166 | -1.68E-16 | 70.69 | 2.51E-10 | 6.53E-10 | 416 | 370 | 88.94% |
| ME10 | 0.3547 | -0.0331 | 3.70E-17 | 34.90 | 1.45E-07 | 3.15E-07 | 240 | 229 | 95.42% |
| ME3 | -0.1789 | -0.2734 | 3.47E-17 | 19.56 | 1.29E-05 | 2.40E-05 | 593 | 560 | 94.44% |
| ME8 | 0.1741 | 0.2650 | 3.24E-17 | 16.67 | 3.86E-05 | 6.28E-05 | 335 | 328 | 97.91% |
| ME11 | -0.3001 | 0.0301 | -6.09E-17 | 13.20 | 1.70E-04 | 2.45E-04 | 173 | 154 | 89.02% |
| ME7 | 0.2566 | -0.1213 | 9.25E-17 | 10.27 | 7.01E-04 | 9.11E-04 | 411 | 409 | 99.51% |
| ME5 | -0.1739 | -0.1885 | 8.10E-18 | 7.17 | 3.98E-03 | 4.71E-03 | 525 | 508 | 96.76% |
| ME12 | 0.0225 | 0.1101 | -1.94E-16 | 0.90 | 0.42 | 0.45 | 108 | 107 | 99.07% |
| ME0 | 0.0233 | 0.0150 | 4.17E-16 | 0.05 | 0.95 | 0.95 | 32 | 0 | 0.00% |

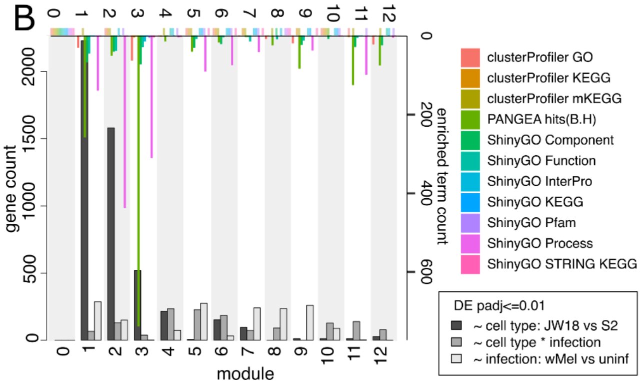


**Figure S9**. *D. melanogaster* DE and WGCNA hits overlap, but not all modules contain terms enriched in known functions. A) Table containing each eigengene module's linear fit with the experimental condition interaction model: ~ cell type + infection state + cell type * infection state, and the proportion of module genes that were significantly differentially expressed. B) Counts of significantly differentially expressed genes per cluster (bottom x-axis and left y-axis). Counts of enriched GO terms and KEGG pathways per *D. melanogaster* module (top x-axis and right y-axis).

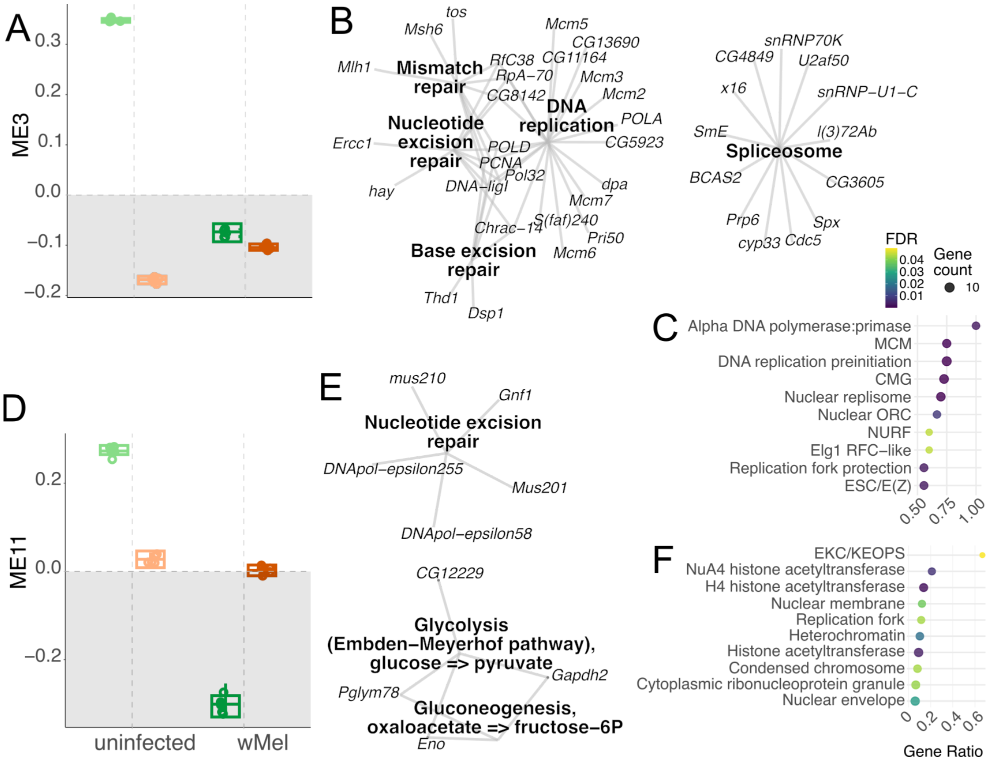


**Figure S10**. *D. melanogaster* WGCNA modules 3 and 11 are associated with uninfected JW18 cells. Functional evidence for eigengene A-C) module 3 and D-F) module 11. A,D) Eigengene value plots for each module. B,E) KEGG enrichment network plots for the genes in each module (FDR-adjusted p-value<0.05). C,F) GO component enrichment plots for each module, plotting the top 5-10 terms by gene ratio (FDR-adjusted p-value<0.05). Key: FDR p-value gradient and gene count dot size for GO plots.

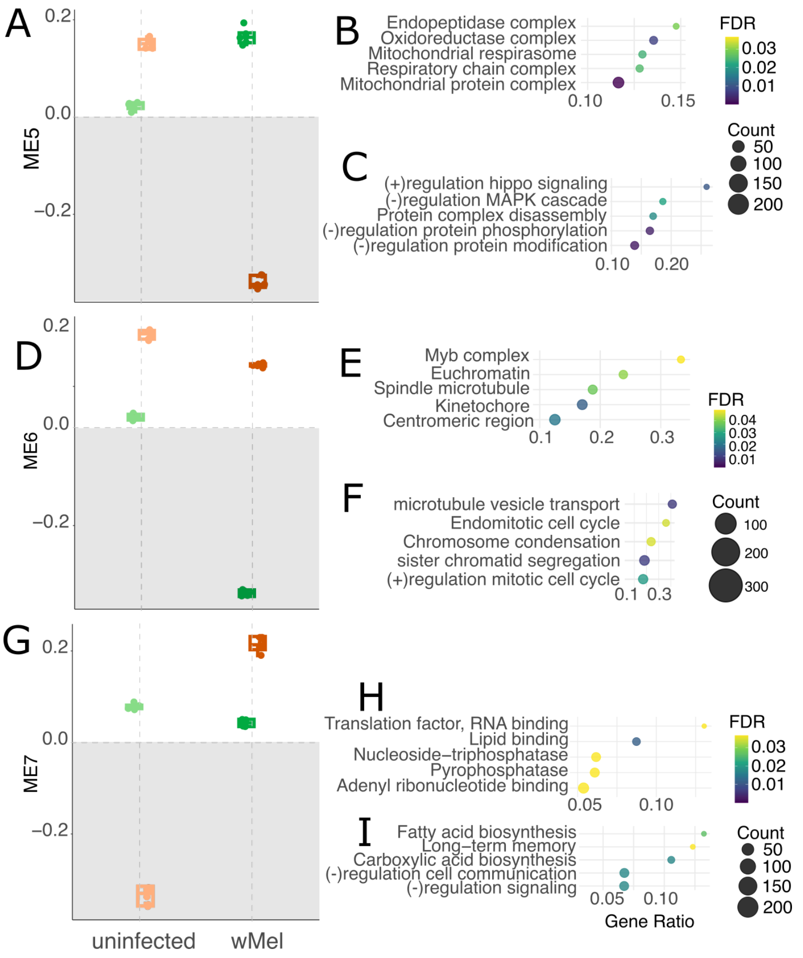


**Figure S11**. *D. melanogaster* WGCNA modules 5, 6, and 7 are associated with complex interactions between cell type and infection state. A,D,G) Eigengene value plots for *D. melanogaster* WGCNA modules. Gene ontology (GO) B,E) component, H) function, and C,F,I) process enrichment plots for each module, plotting the top 5-10 terms by gene ratio (FDR-adjusted p-value<0.05). Keys: FDR p-value gradient and gene count dot size for GO plots.

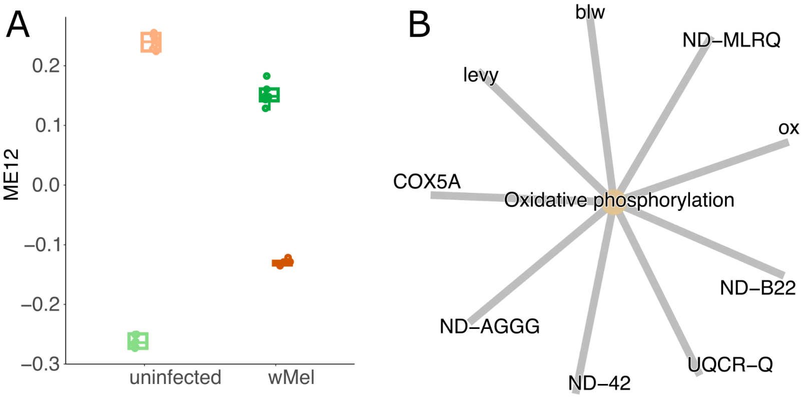


**Figure S12**. *D. melanogaster* constitutive mitochondrial WGCNA module 12 (FDR-adjusted p-value<0.05).

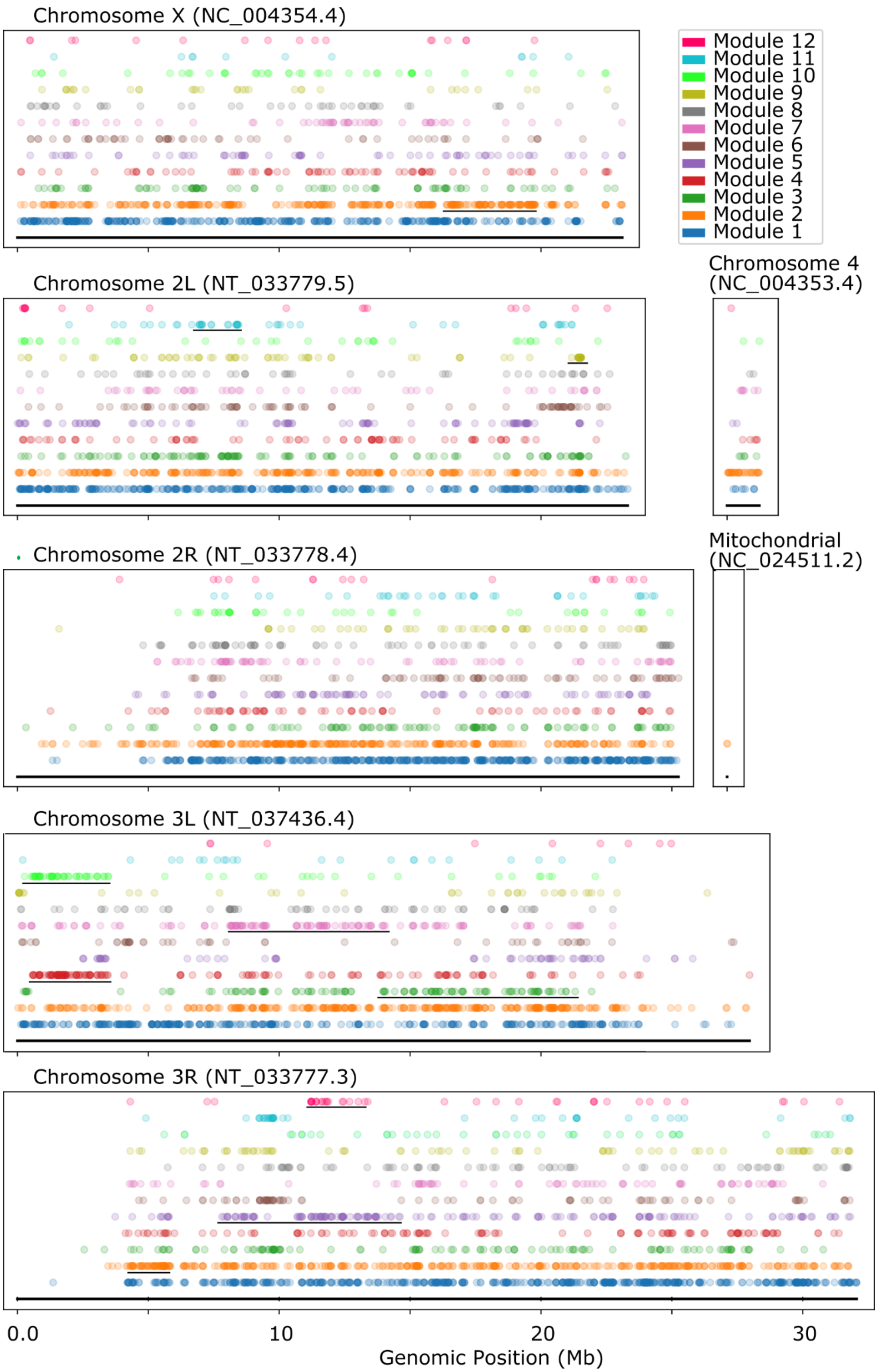


**Figure S13**. Genes comprising several eigengene modules were enriched along particular regions of the *D. melanogaster* genome ($p \leq 0.05$, hypergeometric test; Fig 4P). Black lines indicate regions where module genes are statistically enriched, compared to the background gene density. Genes in JW18-associated module 2 were significantly enriched on the X chromosome and chromosomal arm 3R. JW18 uninfected module 3 and 11 genes were clustered along chromosome arms 3L and 2L, respectively. The modules enriched in infection-associated DE genes, modules 5, 7, and 9, contained genes that were localized along chromosomal arms 3R, 3L, and 2L, respectively. Modules 4 and 10, were enriched in genes differentially expressed due to the interaction between cell type and infection and were both localized along the proximal region of chromosomal arm 3L. The mitochondrial-associated module 12 genes were also localized along the arm of chromosome 3R. These non-random localization patterns of eigengene module genes along the *D. melanogaster* chromosomes, along with the abundance of chromatin-modification hits among the module GO terms suggest that *w*Mel infection is altering host chromatin packaging.

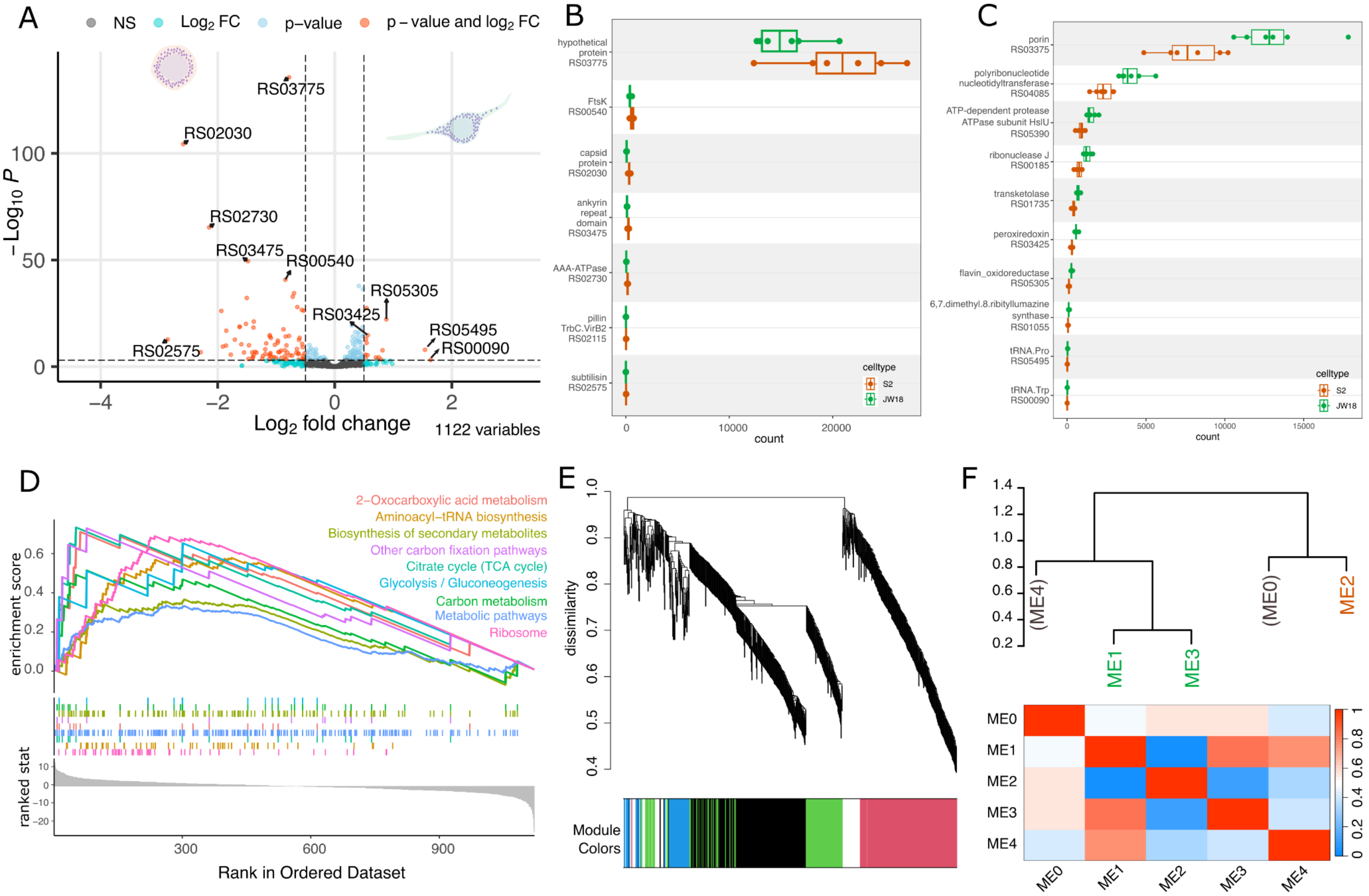


G

| module | logFC | AveExpr | t | P.Value | adj.P.Val | B | genes in module | DE padj>=0.01 genes in module | %DE padj>=0.01 genes in module |
|---|---|---|---|---|---|---|---|---|---|
| ME1 | -0.5717 | -1.85E-17 | -19.51 | 5.01E-10 | 2.50E-09 | 13.558 | 237 | 116 | 48.95% |
| ME2 | 0.5695 | -1.90E-16 | 17.32 | 1.83E-09 | 4.58E-09 | 12.203 | 216 | 106 | 49.07% |
| ME3 | -0.4135 | 2.36E-16 | -3.41 | 0.00567 | 0.00944 | -3.591 | 138 | 30 | 21.74% |
| ME4 | -0.2763 | 8.33E-16 | -1.82 | 0.09602 | 0.12003 | -6.354 | 63 | 1 | 1.59% |
| ME0 | 0.0457 | 3.31E-16 | 0.26 | 0.79581 | 0.79581 | -7.889 | 109 | 0 | 0.00% |

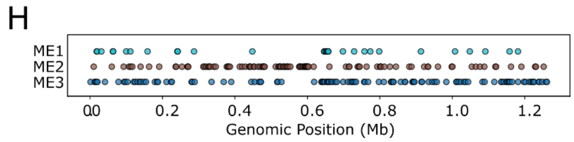


**Figure S14.** *Wolbachia* bulk transcriptome DESeq2 differential expression analysis, GSEA, and WGCNA results. A) Volcano plot of DESeq2 differential expression results. B,C) Transcriptomic read counts for *w*Mel genes highlighted in (A), which are upregulated in B) S2 and C) JW18 cells. D) GSEA, ranking on the "stat" value, showing GO pathways significantly up- and downregulated in JW18 cells relative to S2 cells ($p \le 0.05$). E-F) Weighted gene co-expression network analysis (WGCNA) reveals expression modules associated with cell type, infection, and joint-cell type and infection states. E) Dendrogram and F) heatmap depicting relationships among the modules' eigengenes, the collapsed, combined, and normalized expression of the genes that make up each module. G) Table of module association with the experimental condition interaction model: ~ cell type, and overlap with DE hits. H) Localization of Modules 1-3 genes along the *w*Mel chromosome.

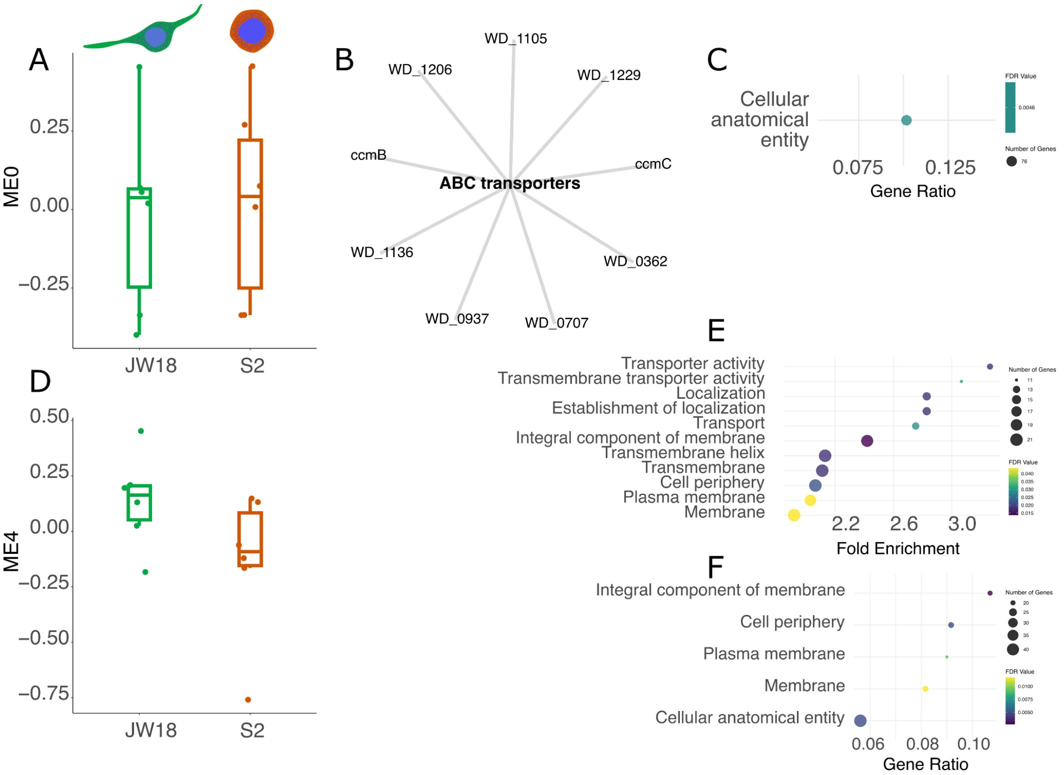


**Figure S15**. wMel constitutive Modules 0 and 4. Eigengene Module 0 A) eigengene values by cell type, B) KEGG enrichment term for genes in the module (FDR-adjusted p-value<0.05), and C) GO component enrichment terms (FDR-adjusted p-value<0.05). Eigengene Module 4 D) eigengene values by cell type and enrichment terms by E) ShinyGO pathway and F) GO component. Keys: FDR p-value gradient and gene count dot size for GO plots.

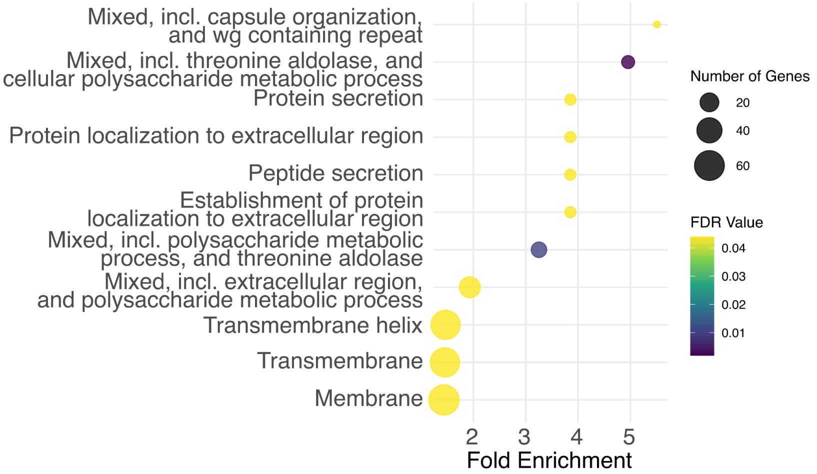


**Figure S16**. *w*Mel Module 2 ShinyGO pathway enrichment terms (FDR-adjusted p-value<0.05). Key: FDR p-value gradient and gene count dot size for GO plots.

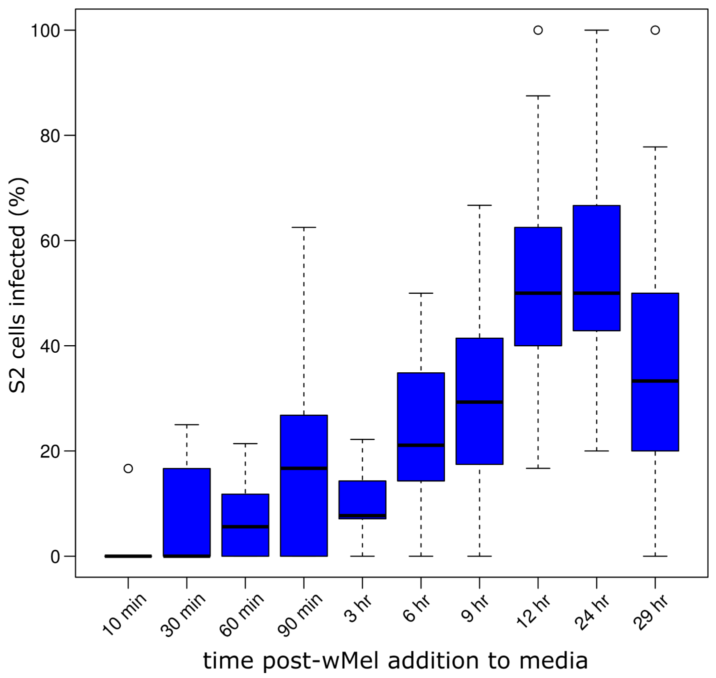


**Figure S17**. Infection rate of S2 cells after cell-free *w*Mel addition to the culture's Shields and Sang M3 medium. FISH staining of *w*Mel's 16S rRNA was used to detect intracellular bacteria. A S2 cell was counted as infected if one or more *w*Mel-FISH puncta was detected.

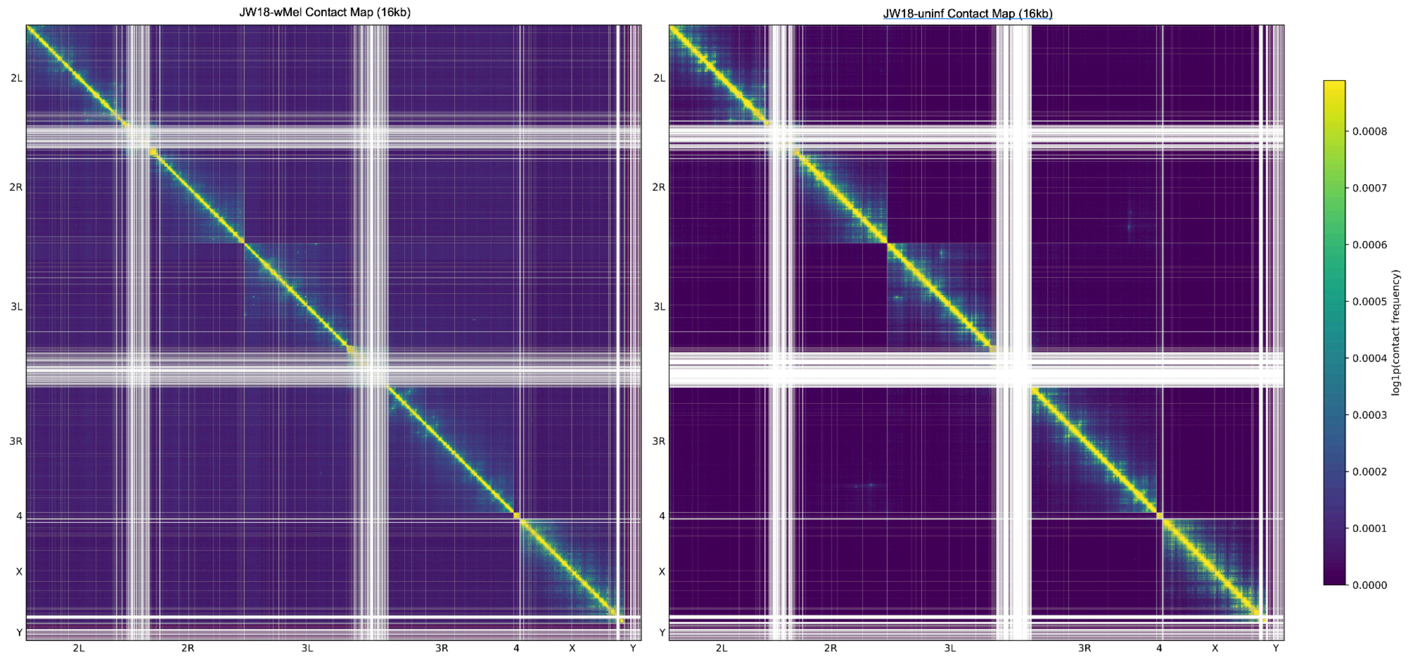


**Figure S18**. *D. melanogaster* Micro-C chromatin contact maps for the JW18 *w*Mel-infected cells (left) and JW18 uninfected cells (right).

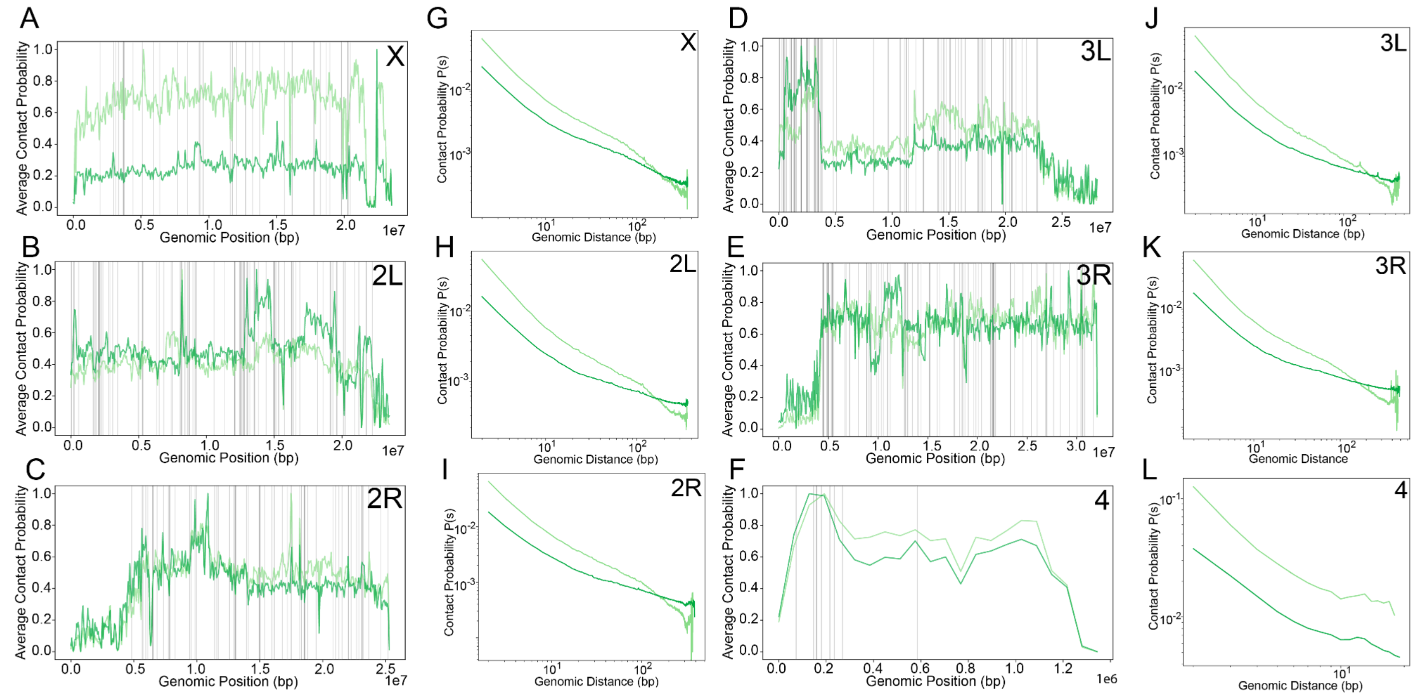


**Figure S19.** Chromatin contact probabilities plotted by *D. melanogaster* chromosome**.** A-F) Average chromatin contact probability plotted along genomic positions for *Drosophila* chromosomes: A) X, B) 2L, C) 2R, D) 3L, E) 3R, and F) 4. Dark green lines represent *w*Mel-infected JW18 cells; light green lines represent uninfected JW18 cells. Grey vertical lines represent locations of topologically associated domains (TADs) differentially detected in *w*Mel-infected and uninfected JW18 cells. G-L) Log-scale plots showing how contact probability decays with increasing genomic distance (in base pairs) for chromosomes: G) X, H) 2L, I) 2R, J) 3L, K) 3R, and L) 4. *w*Mel-infected JW18 cells represented in dark green; uninfected JW18 cells represented in light green.

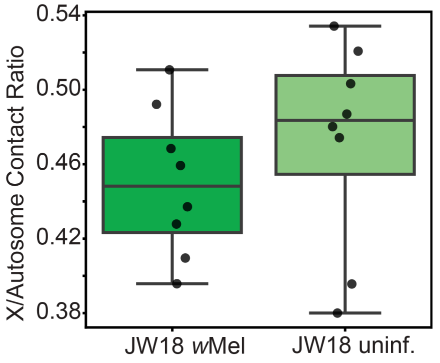


**Figure S20**. X-to-autosome contact ratios per replicate against each autosome (2L, 2R, 3L, 3R). Mann-Whitney U, P = 0.67; n = 2.

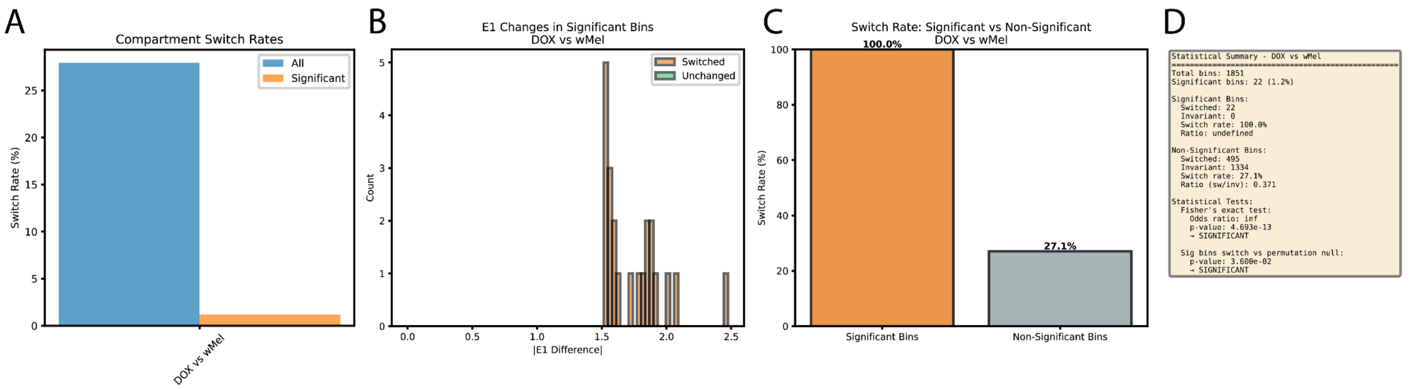


**Figure S21**. Total chromatin compartment switches detected between uninfected and *w*Mel-infected JW18 cells, A) including insignificant compartments. B) Distribution of E1 difference absolute values for the compartments that exhibited significant A/B switches between uninfected and infected conditions. C) Switch rate for compartments in significant and non-significant bins. D) Output from cooltools compartment analysis.

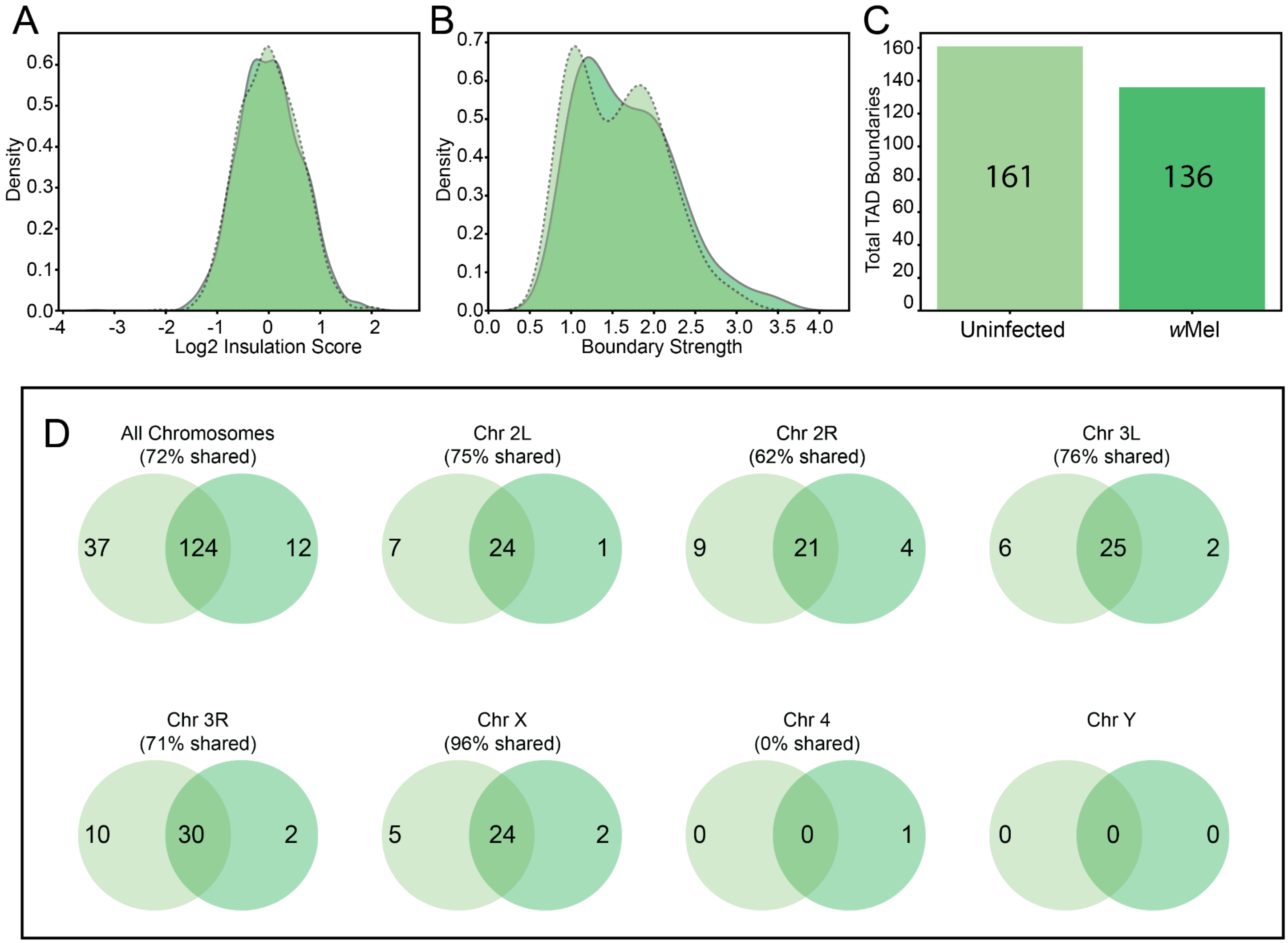


**Figure S22.** Impact of *w*Mel infection on JW18 topologically associating domain (TAD) boundaries. A) Genome-wide distribution of global insulation scores by density for uninfected (dashed light green) and *w*Mel-infected (dark green) conditions. B) Kernel density distribution of TAD boundary strengths across the genome, comparing uninfected (dashed light green) and *w*Mel-infected (dark green) states. C) Total count of TAD boundaries identified in uninfected (light green) and *w*Mel-infected (dark green) conditions. D) TAD overlap between uninfected and *w*Mel-infected conditions.

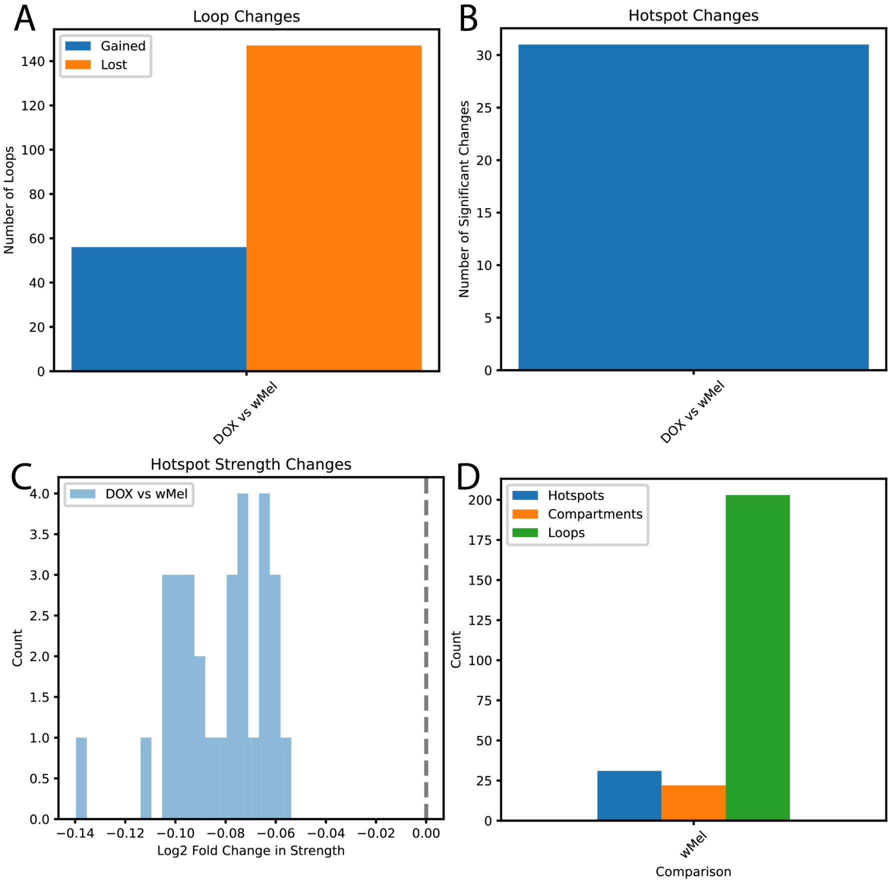


**Figure S23.** Differential contact loop and hotspot analysis. A) Number of hotspots that change between uninfected (DOX) and *w*Mel-infected conditions. B) Direction of differential contact changes within hotspots. C) Barplot of loops gained and lost when cells become stably infected with *w*Mel. D) Summary of significant changes by category.

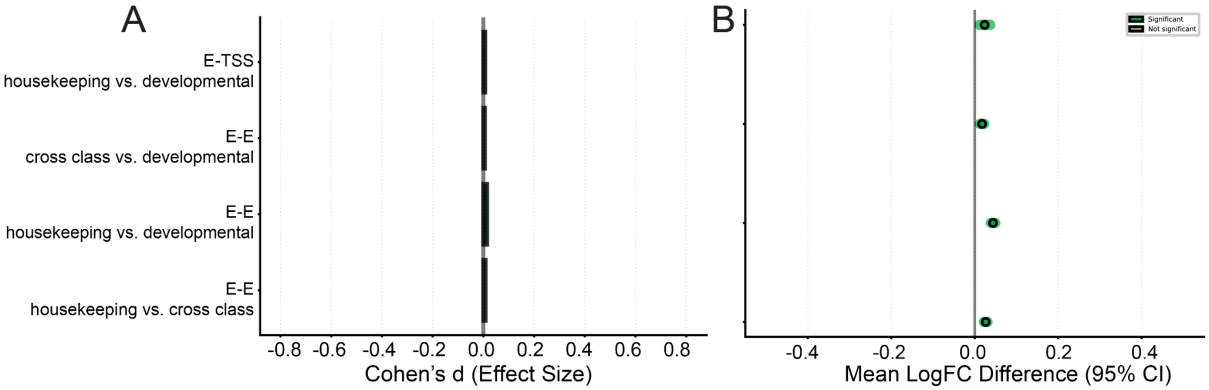


**Figure S24.** Enhancer-promoter enrichment.

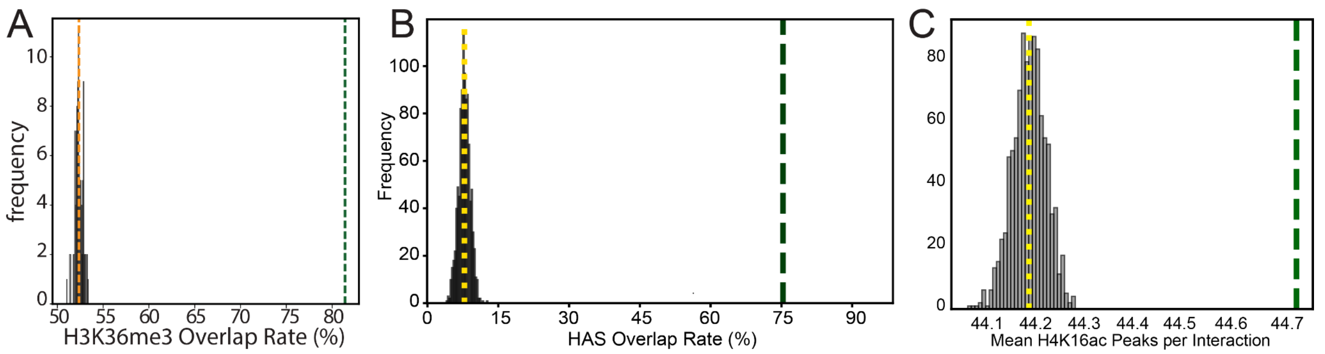


**Figure S25**. Observed differential contact overlap rate with A) H3K36me3 peaks, B) HAS sites, and C) H4K16ac peaks (dashed lines) compared to randomly permuted distributions (grey bars, n=100). Mann-Whitney U test, *** p<0.001.

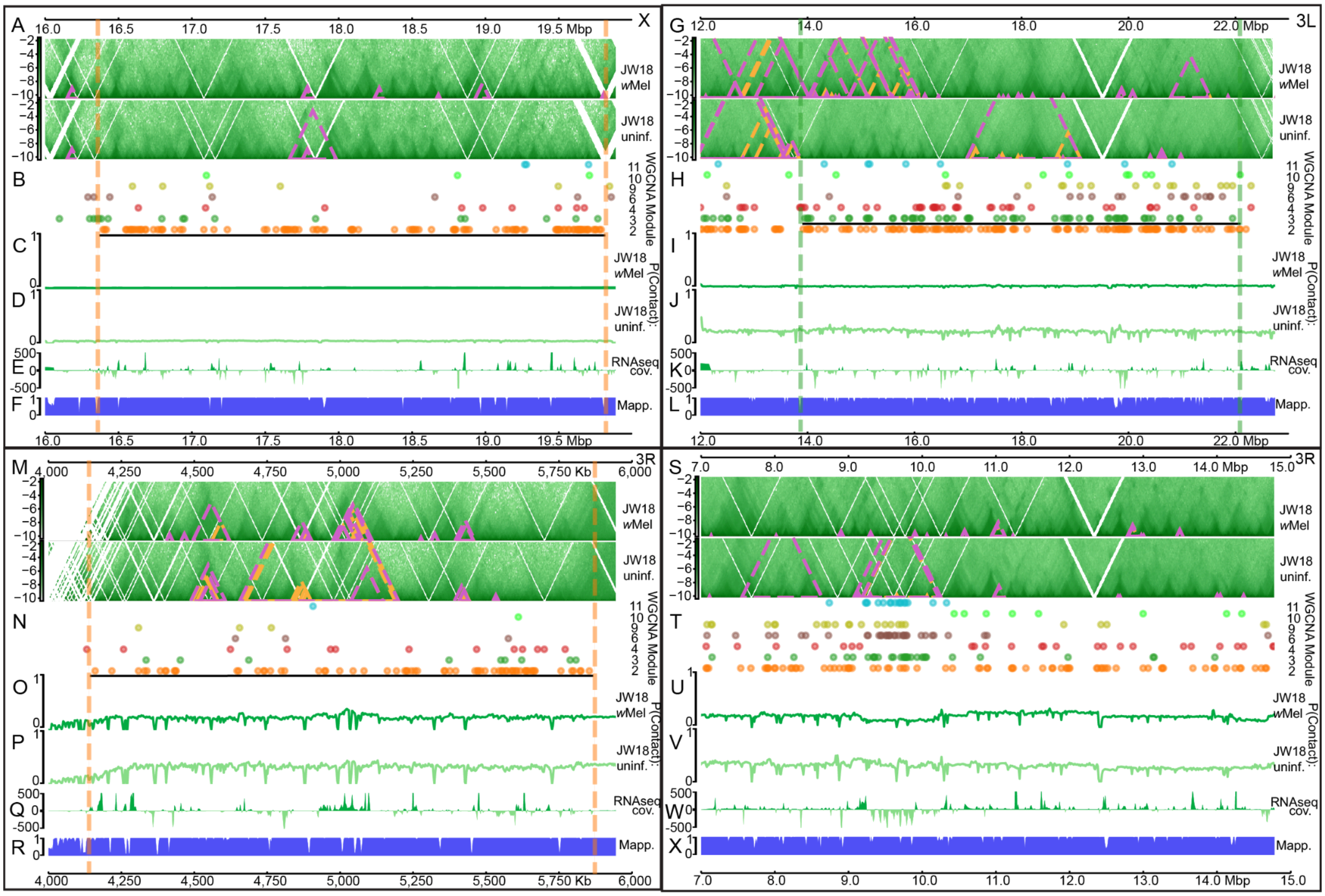


**Figure S25**. Genomic browser views of the Micro-C results for four representative WGCNA module co-localized regions showing multiple data tracks. Regions shown: A-F) Chromosome X:16.0-20.0Mb G-L) Chromosome 3L:12.0-23.0 Mb M-R) Chromosome 3R 4.0-6.0 Mb, S-X) Chromosome 3R 7.0-15.0 Mb. A,G,M,S) Top two tracks: Chromatin contact heatmaps (green intensity) for JW18 *w*Mel-infected and JW18 uninfected cells. Orange outlines highlight significant (q < 0.1) topologically associated domains (TADs), pink outlines indicate significant contact differences (q < 0.05) between uninfected and *w*Mel-infected JW18 cells. B,H,N,T) Middle tracks: Colored dots representing genes from identified eigengene modules. Black lines indicate regions where module genes are statistically enriched, compared to the background gene density.

Module-colored vertical dashed lines mark the boundaries of enriched regions. Next two tracks: Contact probability differences between C,I,O,U) infected (light green) and D,J,P,V) uninfected cells (dark green). E,K,Q,W) RNA-seq coverage differences; (light green representing enriched in uninfected, dark green representing enriched in infected). F,L,R,X) Bottom track: *D. melanogaster* genome mappability (blue).
